# Patterns of Brain Activation and Hippocampal Functional Connectivity Supporting Verbal Memory in Midlife Women

**DOI:** 10.1101/2025.10.13.681463

**Authors:** Katrina A. Wugalter, Rebecca C. Thurston, Minjie Wu, Rachel A. Schroeder, Howard J. Aizenstein, Pauline M. Maki

## Abstract

Women show declines in verbal memory across the menopause transition that may persist into the postmenopause. The goal of the present study was to characterize the patterns of brain activity and hippocampal functional connectivity that support verbal memory performance in midlife postmenopausal women. The study sample included 171 midlife postmenopausal women from the MsBrain I study (mean age = 59.3 years, mean education= 15.7 years, 87.7% white). All participants were cognitively normal, native English speakers, not taking menopausal hormone therapy. Participants completed neuropsychological (California Verbal Learning Test [CVLT]) and neuroimaging assessments, including an fMRI task of verbal encoding and recognition. Findings indicated that during verbal encoding, greater activation of bilateral prefrontal and medial temporal regions, as well as the precuneus, cuneus, caudate, and cerebellar regions, was associated with better performance on CVLT measures, including learning, short- and long-delay recall, and semantic clustering. Functional connectivity from both hippocampi to primarily right prefrontal regions during verbal encoding associated with better CVLT performance. In-scanner word recognition accuracy was more strongly associated with activation of parietal and occipital regions, and with functional connectivity between the right hippocampus and bilateral parietal and temporal regions. Our findings characterize the patterns underlying verbal memory abilities in midlife postmenopausal women. The patterns identified here may act as a foundation for better interpreting the effects of hormonal changes and menopausal symptoms on cognition at midlife, and for identifying neural targets for pharmacological and lifestyle interventions aimed at sustaining women’s memory function.

## 1. INTRODUCTION

Forgetfulness is a common concern among peri- and postmenopausal women (Gold et al., 2000). This concern is substantiated by evidence that neurological changes at menopause may impact cognition (Brinton et al., 2015). Amongst several cognitive domains, verbal memory performance in particular decreases across the menopause transition (Epperson et al., 2013; Greendale et al., 2009; Kilpi et al., 2020; Maki et al., 2021). In large-scale longitudinal studies assessing verbal memory performance with list-learning and story-learning tasks, women perform worse on immediate and delayed recall measures as they transition from the premenopause to perimenopause (Epperson et al., 2013; Greendale et al., 2009; Kilpi et al., 2020; Maki et al., 2021). While some evidence indicates that the decline in verbal learning and/or memory extends into the postmenopause (Epperson et al., 2013; Maki et al., 2021), other evidence suggests that cognitive changes resolve in the postmenopause (Greendale et al., 2009). Understanding the factors that underlie these changes in memory performance at menopause requires a foundational knowledge of typical brain activity and connectivity, but little is known about the patterns of brain functioning that sustain verbal memory abilities in midlife women.

One approach to addressing that gap in knowledge is to examine associations between episodic memory performance and brain function. Very few neuroimaging studies have examined these associations in cognitively normal healthy adults at midlife (e.g., Grady et al., 2006, Jacobs et al., 2016; Kwon et al., 2016; Schroeder et al., 2024). Two of those studies focused on women and each examined those associations as secondary analyses to help interpret a limited set of menopause- and/or estrogen-related findings (Jacobs et al., 2016; Schroeder et al., 2024). In a cross-sectional study of peri-, pre-, and postmenopausal women and midlife men, activity of the left hippocampus during a verbal encoding task decreased across the menopause transition, while bilateral hippocampal connectivity increased (Jacobs et al., 2016). Low levels of estradiol (E2) were associated with greater left-right hippocampal functional connectivity. In addition, postmenopausal women who performed well on an associative episodic memory task (i.e., the Face-Name Associative Memory task) showed less bilateral hippocampal connectivity and more recruitment of the prefrontal cortex (PFC) during verbal encoding relative to middle and low performing women (Jacobs et al., 2016). In a sample of late midlife postmenopausal women, we found that E2 was positively associated with bilateral (primarily left) frontal and temporal activation, while estrone (E1) was positively associated with a wider array of regions including bilateral temporal and right frontal regions (Schroeder et al., 2024). Notably, activation in key regions positively associated with E2 (i.e., left inferior and middle frontal gyri) was also positively associated with measures of verbal learning and memory (Schroeder et al., 2024). These initial studies suggest that activation of frontal and temporal regions, as well as strong PFC-hippocampus connectivity in the left hemisphere may be critical for maintaining verbal memory encoding in midlife postmenopausal women.

To more directly establish the brain circuitry supporting successful verbal memory performance, it is necessary to comprehensively examine the patterns of brain activation and connectivity associated with performance on validated, sensitive measures of verbal episodic memory. Our understanding of the effects of hormonal changes, menopausal symptoms, and Alzheimer’s disease risk factors on women’s cognition relies upon a foundational knowledge of the typical patterns supporting memory performance in midlife women that has yet to be delineated. To address this gap, we identified patterns of brain activation and hippocampal functional connectivity during verbal encoding associated with successful verbal memory abilities in an appreciably large (n=171), well-characterized sample of late midlife postmenopausal women participating in the MsBrain study.

## 2. MATERIALS AND METHODS

### 2.1. Participants

The study sample comprised women from the MsBrain study, a cohort study of brain health, menopause, and vasomotor symptoms (VMS; hot flashes, night sweats) in midlife women initiated in 2017 in Pittsburgh, Pennsylvania, USA (Thurston et al., 2023). Participants in MsBrain (N = 274) were recruited from two sources: 170 participants partook in MsHeart, a previous study of VMS and cardiovascular health at the University of Pittsburgh from 2012-2015 (Thurston et al., 2016), and 104 participants were recruited from the Pittsburgh community. Full inclusion and exclusion criteria for the MsHeart and MsBrain studies can be found in prior publications (Schroeder et al., 2024; Thurston et al., 2016; 2023). Participants from MsBrain were included in the present analysis if they: were cognitively normal as determined by their Montreal Cognitive Assessment score (MoCA; Nasreddine et al., 2005), using race- and education-based cut-off scores (Milani et al., 2018); were a native English speaker; had valid neuroimaging data (i.e., task-based activation and functional connectivity, task performance, T1 structural brain imaging for co-registration); and had valid California Verbal Learning Test (CVLT) data, resulting in N = 193. Of these 193 eligible participants, an additional five were excluded due to incidental MRI findings (e.g., mass or vascular formation identified on scan), five due to excessive motion in the scanner, nine due to cognitive or neurological issues (e.g., tardive dyskinesia, previous experience with neuropsychological battery, recent chemotherapy, CVLT performance more than 3 SD below the mean), and three due to failure to perform above chance on the in-scanner functional magnetic resonance imaging (fMRI) task, resulting in a final analytic sample of 171 women.

### 2.2. Experimental Design

The MsBrain study procedures included screening procedures, a medical history, neuroimaging, and neuropsychological testing. This study analyzed the verbal memory task-based neuroimaging and select neuropsychological assessments. The University of Pittsburgh Human Research Protection Office approved the research procedures, and all participants provided informed consent.

#### 2.2.1 Neuropsychological Measures

##### 2.2.1.1 Montreal Cognitive Assessment

The participants completed the Montreal Cognitive Assessment (MoCA) to assess global cognitive ability (Nasreddine et al., 2005). The MoCA is a 10-minute screening tool to detect possible mild cognitive impairment (MCI) or mild Alzheimer’s dementia. A score < 26 out of 30 points on the MoCA indicates possible MCI, with differing cutoffs varying by race and education (see Milani et al., 2018). Only participants considered cognitively normal were included in the present analyses.

##### 2.2.1.2. California Verbal Learning Test

Participants completed the California Verbal Learning Test, Version 2 (CVLT), a sensitive list-learning assessment of verbal episodic learning and memory (Delis et al., 2002). Declines in list-learning measures occur with natural aging, but accelerated declines are predictive of the later development of Alzheimer’s dementia and impairment on list- and story-learning measures is used in the diagnosis of Alzheimer’s disease (Balthazar et al., 2010; Estevez-Gonzalez et al., 2003; Lange et al., 2002). During administration of the CVLT, study staff reads aloud a 16-item word list containing nouns from four semantic categories (i.e., vegetables, desserts, sports equipment, office supplies). The words are in a fixed order, repeated each trial for five trials, with no consecutive words from the same category. After each trial, participants recall as many words as possible. After the list is read five times, an interference list containing unique words is presented. Participants recall as many words as possible from the original list immediately after and 20 minutes after the interference list. The primary outcomes are: total learning (total number of words recalled on trials 1-5); short-delay free recall (number of words recalled immediately after the interference list); long-delay free recall (number of words recalled 20 minutes after the interference list); and semantic clustering. A semantic cluster is at least two consecutively recalled words from the same semantic category (e.g. “cabbage” followed by “broccoli”). Semantic clustering is an effective strategy for episodic encoding which relies on the prefrontal cortex (Fletcher et al., 1998; Gabrieli et al., 1998; Savage et al., 2001), shows a female advantage (Kramer et al., 1988), and may be influenced by sex hormones as evidenced by greater semantic clustering abilities among menopausal hormone therapy (MHT) users compared to non-users (Maki et al., 2001; Resnick et al., 1998). Individual weighted clustering scores were calculated for learning, short-delay, and long-delay trials by assigning 1 point for each 2-word cluster, 2 points for each 3-word cluster, and 3 points for each 4-word cluster. The final measure of semantic clustering used in the present study was a combined standardized score of the individual weighted clustering scores for the learning, short- and long-delay trials.

#### 2.2.2 Neuroimaging

Participants underwent neuroimaging at the MR Research Center of the University of Pittsburgh with a 3T Siemens Tim Trio MRI scanner and a Siemens 64- channel head coil. T1-weighted structural images were acquired in the axial plane using a magnetization-prepared rapid gradient echo sequence (T1w MPRAGE) with the following parameters: TR =□2400 ms, TE = 2.22 ms, flip angle□= 8°, FOV□=□240 × 256 mm^2^, matrix = 300 × 320, slice thickness/gap□=□0.8/0 mm (voxel size□=□0.8□× 0.8 × 0.8 mm^3^), and 208 slices. T1w MPRAGE images were used to facilitate the normalization of fMRI data into the Montreal Neurological Institute (MNI) template. Whole-brain fMRI data were acquired axially (i.e., parallel to anterior commissure - posterior commissure line) using gradient-echo echo-planar imaging sequence with the following parameters: repetition time =□1 second, echo time =□30 ms, flip angle□=□45°, FOV□= 220 × 220 mm2, acquisition matrix 96 × 96, slice thickness/gap = 2.3/0 mm (voxel size = 2.3 × 2.3 × 2.3 mm^3^), 60 axial slices.

##### 2.2.2.1. In-Scanner Verbal Memory Task

While in the scanner, participants completed a verbal memory task comprising verbal encoding and recognition tasks modeled after the Hopkins Verbal Learning Test (Brandt & Benedict, 2001) and the CVLT (Delis et al., 2002). During the encoding task, participants were presented with one noun on the screen at a time, asked to press the button after seeing each word, and instructed to try to remember all the words for a later memory test. The encoding task included five experimental blocks alternating with five control blocks, for a total of 60 presented words. In the experimental blocks, participants were shown a total of 30 unique nouns comprising 6 categorical exemplars from five semantic categories (i.e., fruits, nature, body parts, insects, and furniture). Each 6-item experimental block included two unique exemplars (e.g., “ladybug,” and “grasshopper” as exemplars of insects) from three categories. In each 6-item control block, participants were presented the same two nouns (i.e., “sailboat” and “ballet”) in repeating order. Each item in the encoding task was presented for 3 seconds, with a 1.5-sec interstimulus interval. Participants completed a practice test prior to entering the scanner.

Fifteen minutes following the encoding task, during which structural images were acquired, participants completed a yes/no recognition task with experimental and control blocks resembling the encoding task. In the experimental blocks, participants were shown a total of 30 words: 15 previously presented category exemplars, 7 related distractor words, and 8 unrelated distractor words. In the control blocks, they were repeatedly shown the words “sailboat” and “ballet”. Each item in the recognition task (60 total words) was presented for 2.7 seconds, with a 0.6-sec interstimulus interval. The primary behavioral outcome was the number of words correctly recognized during the experimental blocks divided by the total number of correct words (i.e., experimental recognition accuracy).

##### 2.2.2.2. Post-Scanner Verbal Free Recall

After the scanning session, participants were asked to freely recall as many words as they could from the in-scanner verbal memory task. The primary outcomes were: Free Recall Total (total number of words recalled from the in-scanner task) and Free Recall Semantic Clustering. A semantic cluster was defined as at least two consecutively recalled words from the same semantic category (e.g., “grasshopper” followed by “ladybug”). The final Free Recall Semantic Clustering score was a standardized weighted clustering score which assigns 1 point for each 2-word cluster, 2 points for each 3-word cluster, and 3 points for each 4-word cluster.

#### 2.2.3. Additional Measures

Demographics were assessed using questionnaires. Age was reported in years. Education was reported as years of completed education. Race/ethnicity was self-reported and later categorized as White vs. Black, Asian or mixed race for analyses.

### 2.3. Statistical Analysis

#### 2.3.1. Neuroimaging Analyses

Blood-oxygen-level-dependent (BOLD) fMRI data from the verbal encoding task was preprocessed in Statistical Parametric Mapping (SPM12) for slice timing correction, motion correction, image co-registration and normalization, and 8-mm Gaussian imaging smoothing.

For brain activation analyses, first-level analyses were conducted using a general linear model (GLM) that included regressors for experimental and control blocks, with six motion parameters as covariates of no interest. Subject-specific contrast images were subsequently generated for the experimental > control blocks ((i.e., novel words minus repeated words).

For functional connectivity analyses, generalized psychophysiological interaction (gPPI) analysis (Friston et al., 1997) was conducted to estimate hippocampal connectivity during the verbal encoding task. Seed regions were defined as the left and right hippocampus based on the Automated Anatomical Labeling (AAL) atlas (Tzourio-Mazoyer et al., 2002). Principal time series (i.e., the beta value) were generated for each seed region, left and right hippocampus, using singular value decomposition (in MATLAB) from hippocampal fMRI data during the verbal encoding task. Principal time series of the seed region (left or right hippocampus), task conditions (experimental and control blocks), interaction variables (seed times series * task condition), as well as motion parameters were included in the design matrix. Functional connectivity maps (left or right hippocampus) during verbal encoding (novel words minus repeated words) were computed for each participant.

For second-level analyses, regression analyses were conducted to test the associations between verbal memory measures (i.e., CVLT total learning, CVLT short- and long-delay recall, CVLT semantic clustering, and in-scanner recognition) with brain activation or hippocampal functional connectivity during verbal encoding, controlling for age and years of education. To control for multiple comparisons in the activation and functional connectivity analyses, joint height and extent thresholds were determined using Monte Carlo simulations (10,000 iterations) with an a priori whole-brain grey matter mask (AlphaSim, AFNI) for a corrected p < 0.05 (Cox, 1996).

#### 2.3.2. Post-Hoc Analyses

All post-hoc statistical analyses were conducted using R Statistical Software (Version 4.3.1; R Core Team, 2023). For post hoc analyses, the mean beta values were extracted from the significant region of interest (ROI) masks from the second-level activation or functional connectivity analyses. Multiple linear regressions were used to evaluate the associations between verbal memory measures and activation or hippocampal functional connectivity (regional beta values) during verbal encoding, controlling for age, race (white, non-white), and years of education.

To compare performance on the CVLT with performance on the fMRI verbal memory task, we ran linear regression models to examine the relationships between the CVLT measures, in-scanner recognition accuracy measure, and free recall measures assessed directly after scanning session, adjusting for age, race, and years of education.

As a supplementary analysis to compare our findings with prior research (i.e., Jacobs et al., 2016), we divided our sample into tertiles by performance on each CVLT measure. Multiple linear regression analyses were conducted to evaluate whether CVLT performance group (low, middle, high) predicted fMRI measures (activation, hippocampal functional connectivity), controlling for age, race, and education. Supplemental tertile analyses were restricted to the inferior frontal gyrus and hippocampal regions, which were selected *a priori* due to evidence of their associations with episodic memory performance (Jacobs et al., 2016).

## 3. RESULTS

### 3.1. Participants

The final sample comprised 171 postmenopausal women (see Table I).

**TABLE I.**
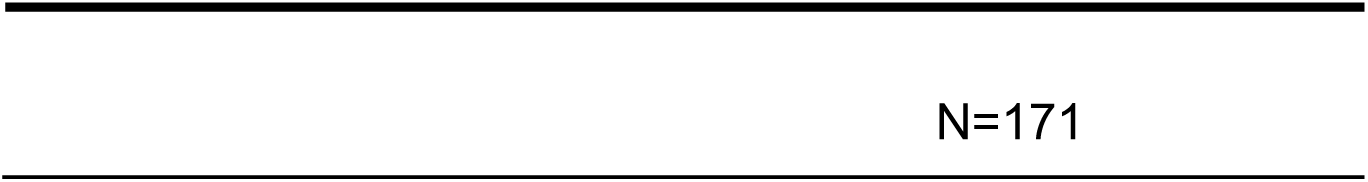
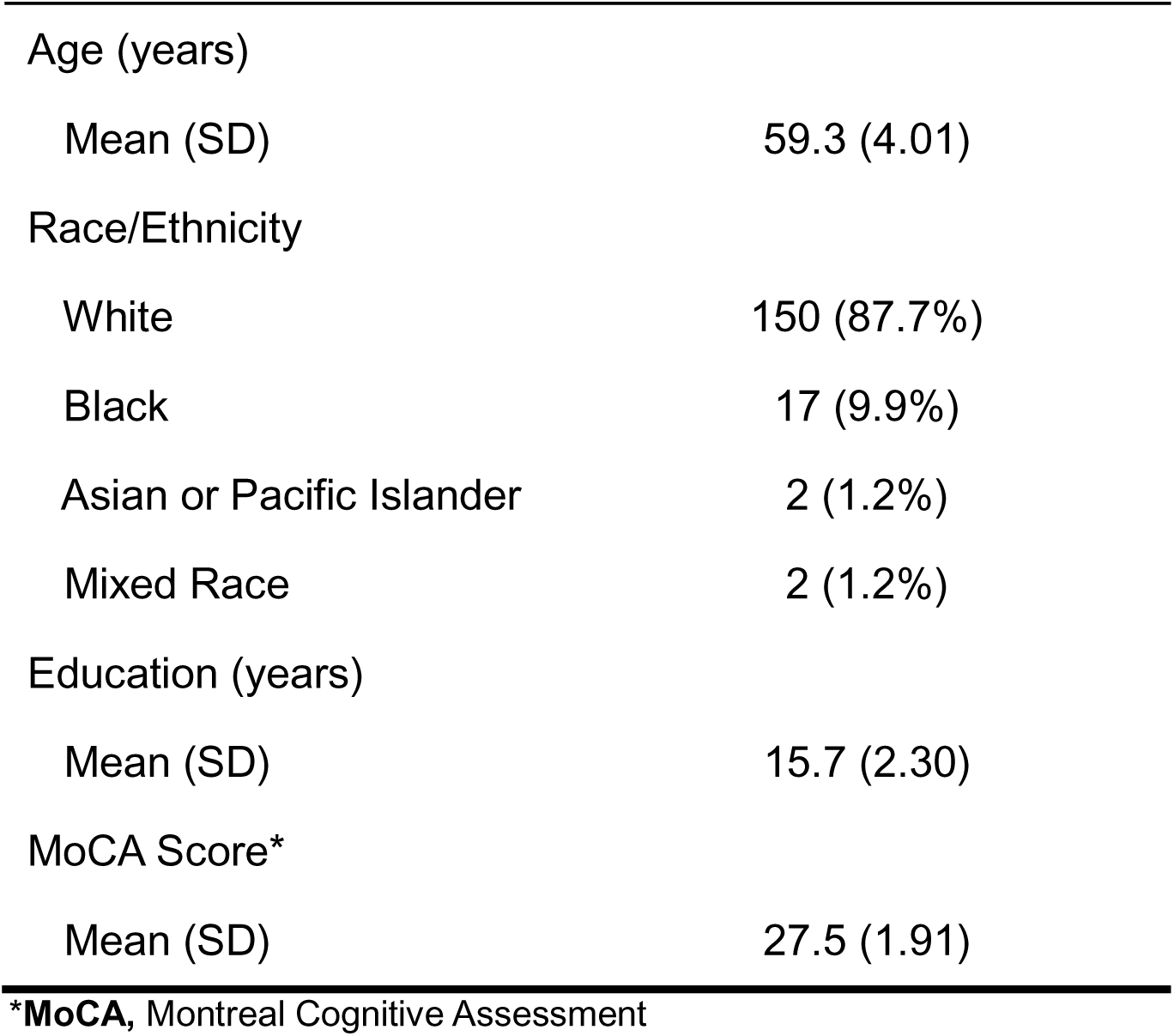
PARTICIPANT DEMOGRAPHICS.

### 3.2. Verbal Memory Performance

On average, participants recalled 54.1 ± 8.19 words across all five learning trials out of a possible 80 words. After a short delay (i.e. immediately after hearing an interference list), participants recalled on average 11.7 ± 2.72 words from the 16-word list. After a long delay (i.e., 20 minutes after hearing an interference list), participants recalled on average 12.5 ± 2.72 words from the 16-word list.

On average, participants’ accuracy on the in-scanner recognition task was 79.2% (SD = 9.4). Experimental accuracy was significantly associated with all CVLT measures, including total learning (*b* = 24.22, *SE* = 6.87, *p =* .0005), short delay recall (*b* = 8.17, *SE* = 2.24, *p* = .0004), long delay recall (*b* = 9.32, *SE* = 2.16, *p* = 2.62 x 10^-5^), and semantic clustering (*b* = 1.92, *SE* = 0.78, *p* = .015).

#### 3.2.1. Correlations between CVLT and Post-Scanner Free Recall Measures

After exiting the fMRI scanner, participants recalled on average 10.03 ± 5.97 words from the 30 target items shown inside the scanner. The total number of freely recalled words from the in-scanner task was associated with CVLT learning (*b =* 0.29, *SE =* 0.047, *p* = 2.6 x 10^-9^), short delay recall (*b* = 0.79, *SE* = 0.144, *p* = 1.67 x 10^-7^), and long delay recall measures (*b* = 0.75, *SE* = 0.148, *p* = 1.11 x 10^-6^). The weighted clustering score after the in-scanner task was associated with semantic clustering on the CVLT (*b* = 0.35, *SE* = 0.074, *p* = 5.02 x 10^-6^).

### 3.3. Activation

#### 3.3.1. CVLT Measures

BOLD activation during the verbal encoding task associated with all CVLT measures of interest in several brain regions (Table II). Total learning score was significantly positively associated with activation of the right cerebellum, right precuneus, left middle frontal gyrus, left inferior temporal gyrus, left superior parietal lobule, left anterior cingulate cortex, and bilateral hippocampi, inferior and medial frontal gyri, caudate, and cuneus (Figure 1). In all significant regions, greater activity was associated with better learning.

**Figure 1.**
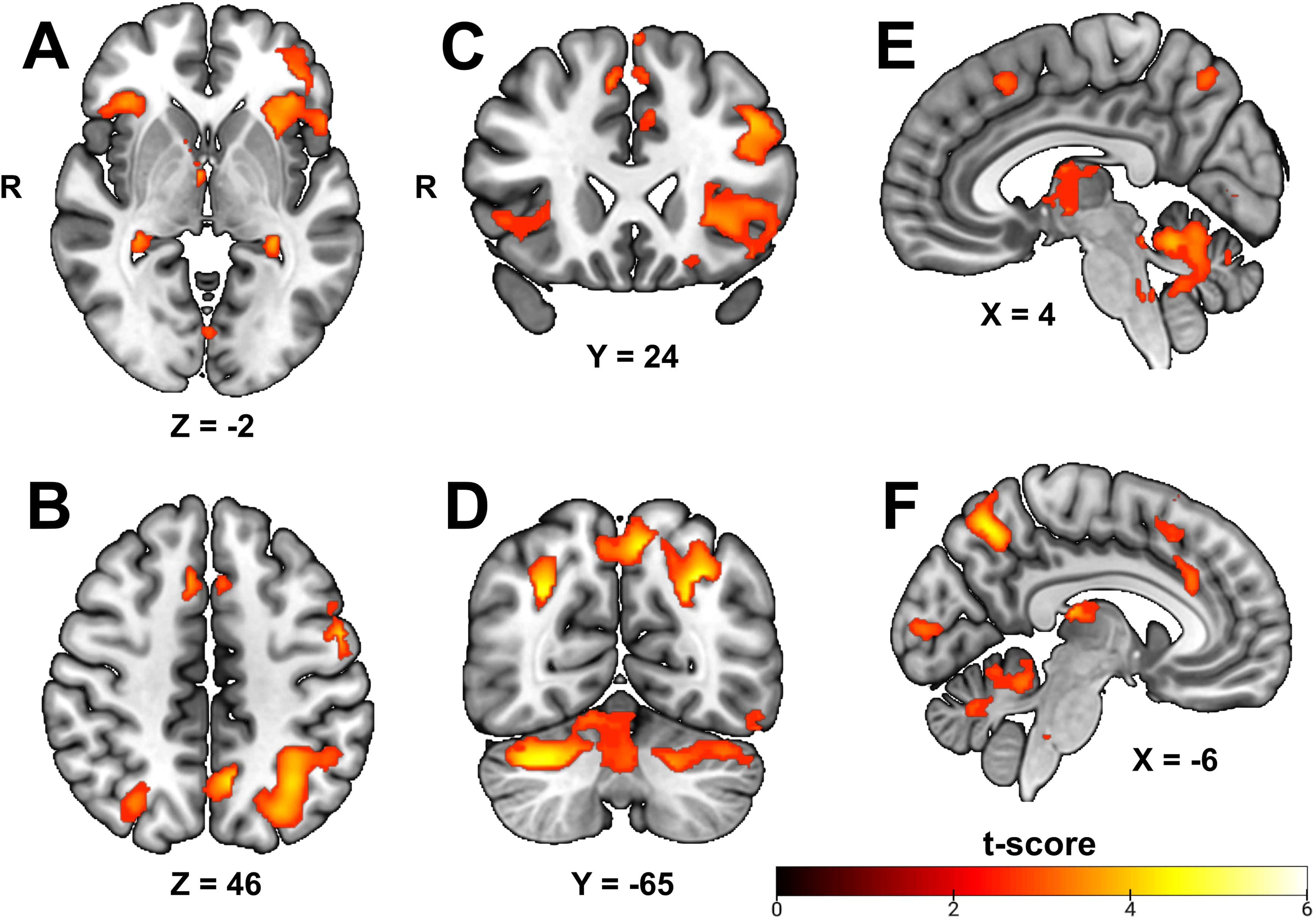
Regional activation positively associated with out-of-scanner total learning score. Functional magnetic resonance imaging (fMRI) results significantly associated with CVLT learning score at p < .05, cluster corrected. For related statistical findings, see Table II. **A,** Inferior axial perspective. **B,** Superior axial perspective. **C,** Anterior coronal perspective. **D,** Posterior coronal perspective, **E,** Right medial sagittal perspective. **F,** Left medial sagittal perspective.

**TABLE II.**
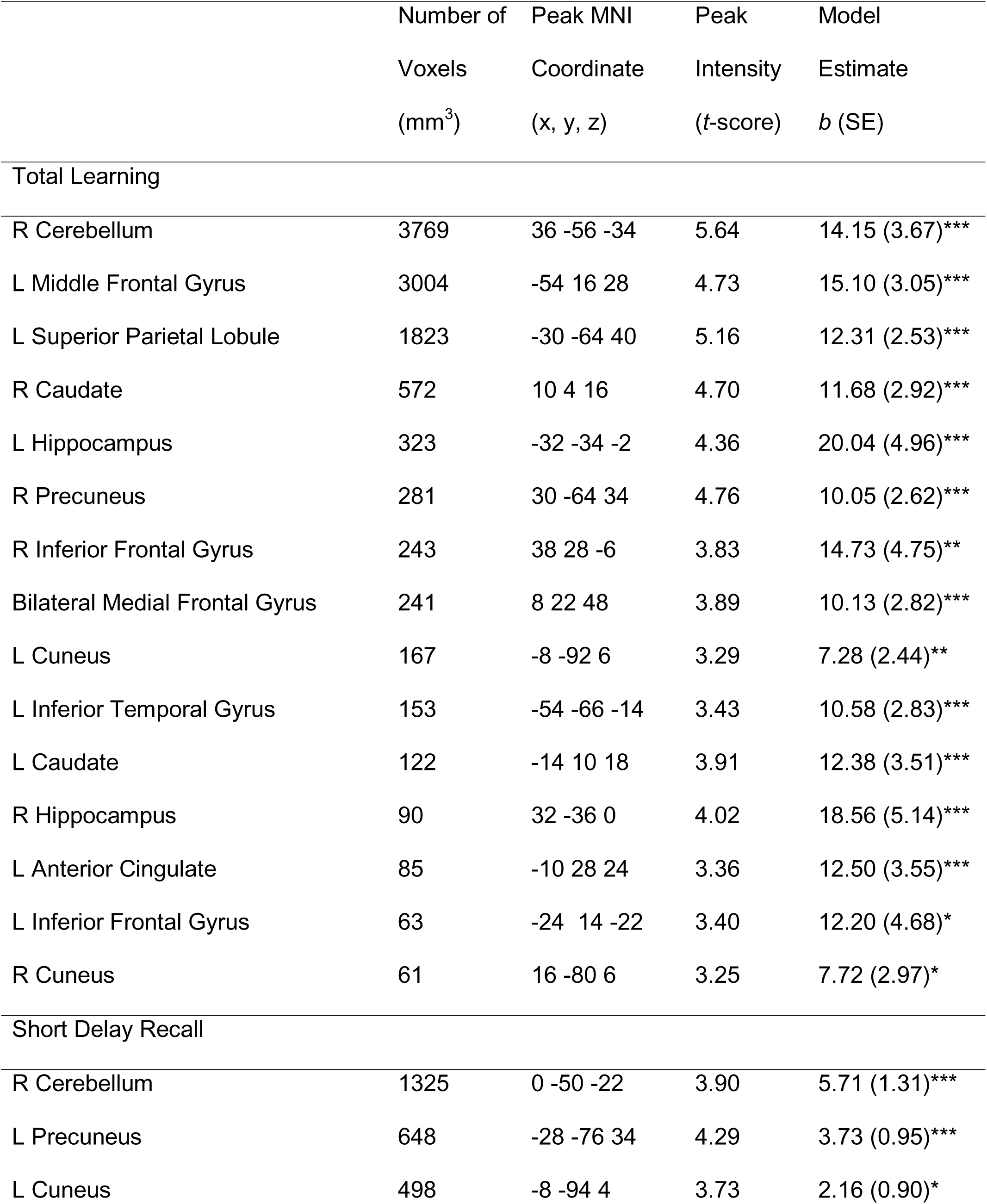
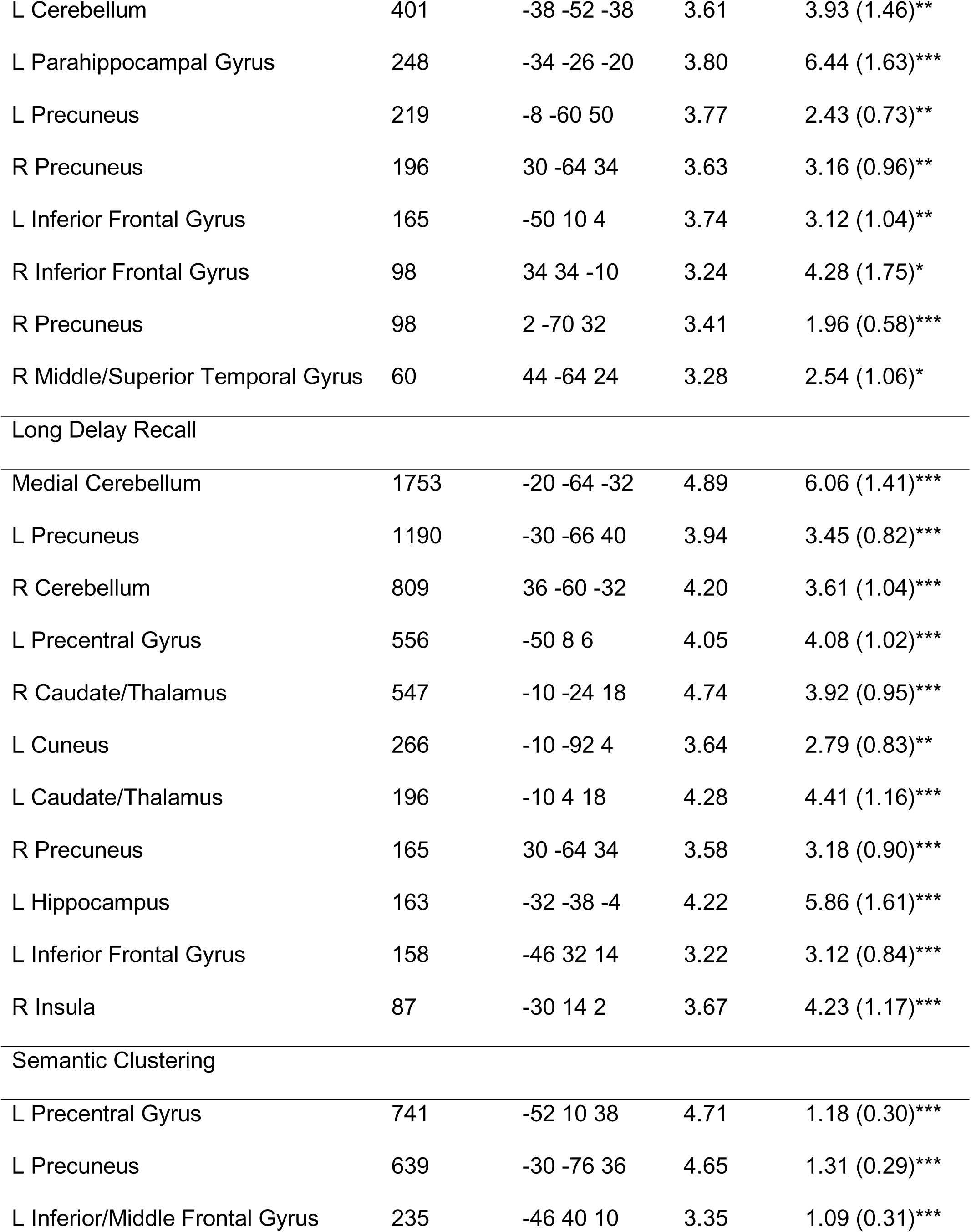
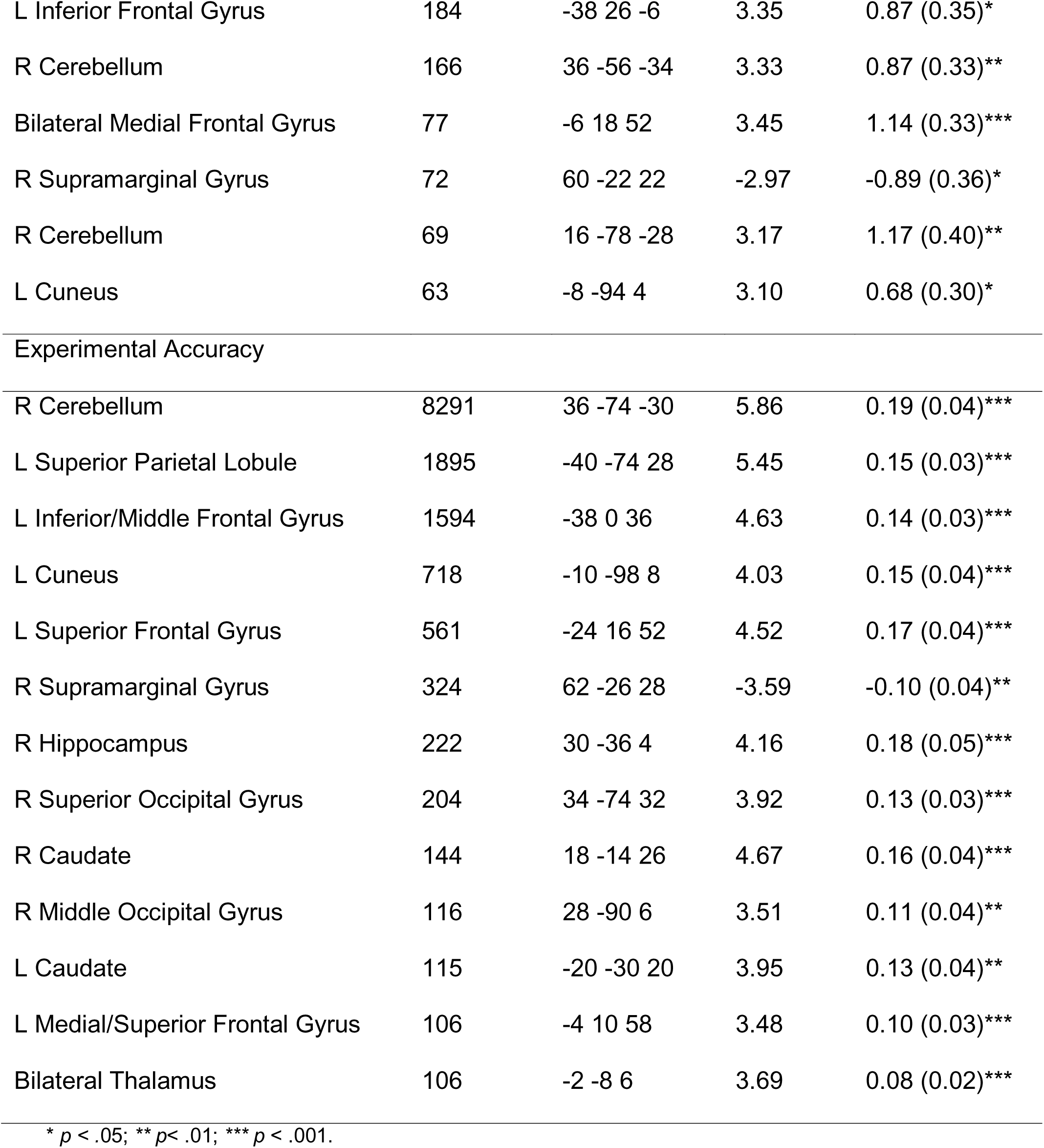
REGIONS OF ACTIVATION ASSOCIATED WITH VERBAL MEMORY MEASURES.

Short delay recall was associated with activity of the right middle/superior temporal gyrus, left cuneus, left parahippocampal gyrus, and bilateral cerebellum, inferior frontal gyrus, and precuneus. In all significant regions, stronger activity was associated with better short delay recall (Figure 2).

**Figure 2.**
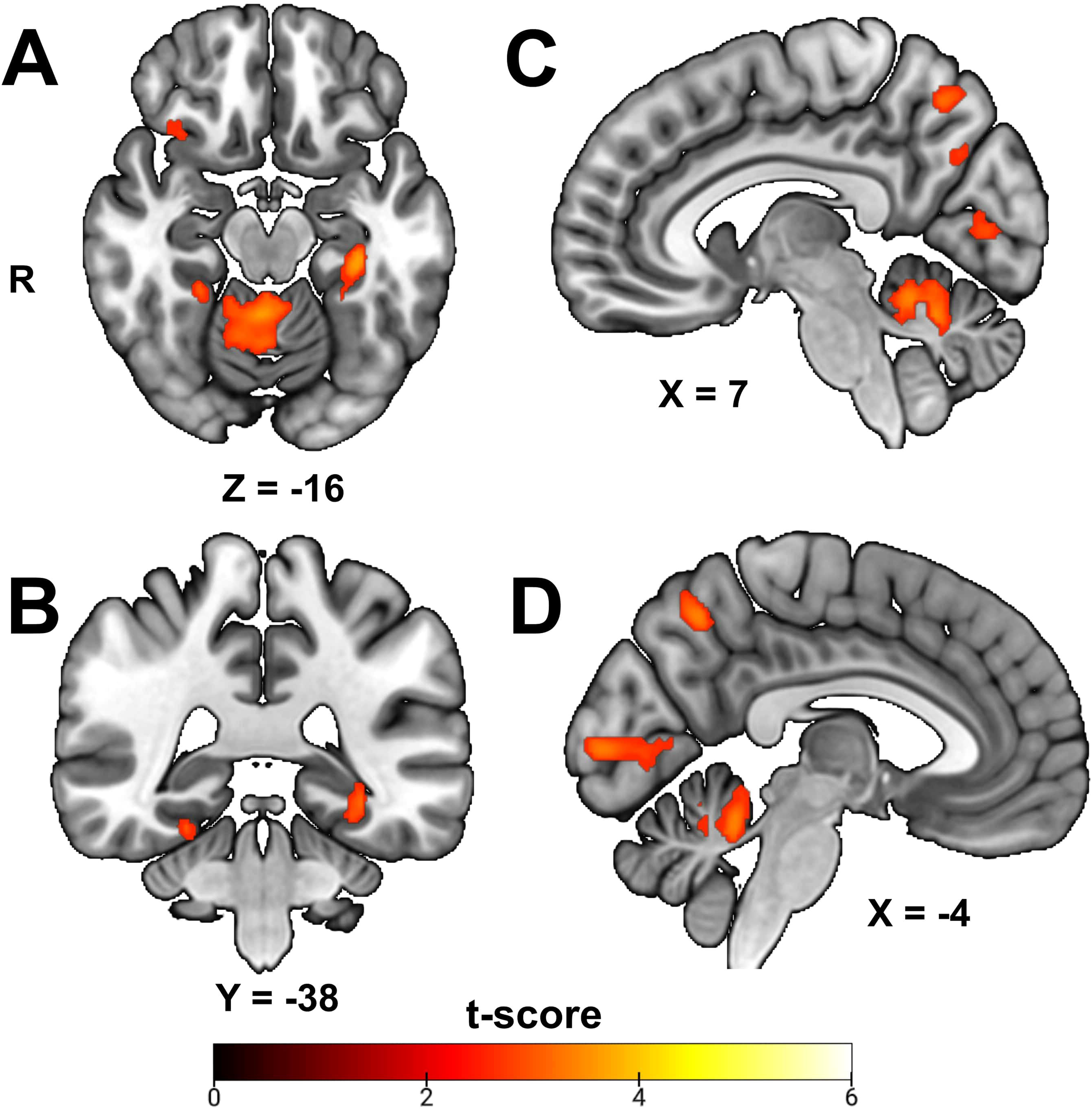
Regional activation positively associated with out-of-scanner short delay recall. Functional magnetic resonance imaging (fMRI) results significantly associated with CVLT short delay recall at p < .05, cluster corrected. For related statistical findings, see Table II. A, Inferior axial perspective. B, Posterior coronal perspective. C, Right medial sagittal perspective. D, Left medial sagittal perspective.

Long delay recall was associated with activity of the right and medial cerebellum, right insula, left hippocampus, left cuneus, left precentral gyrus, left inferior frontal gyrus, and bilateral caudate and precuneus. Greater activity was associated with better long delay recall in all those regions (Figure 3).

**Figure 3.**
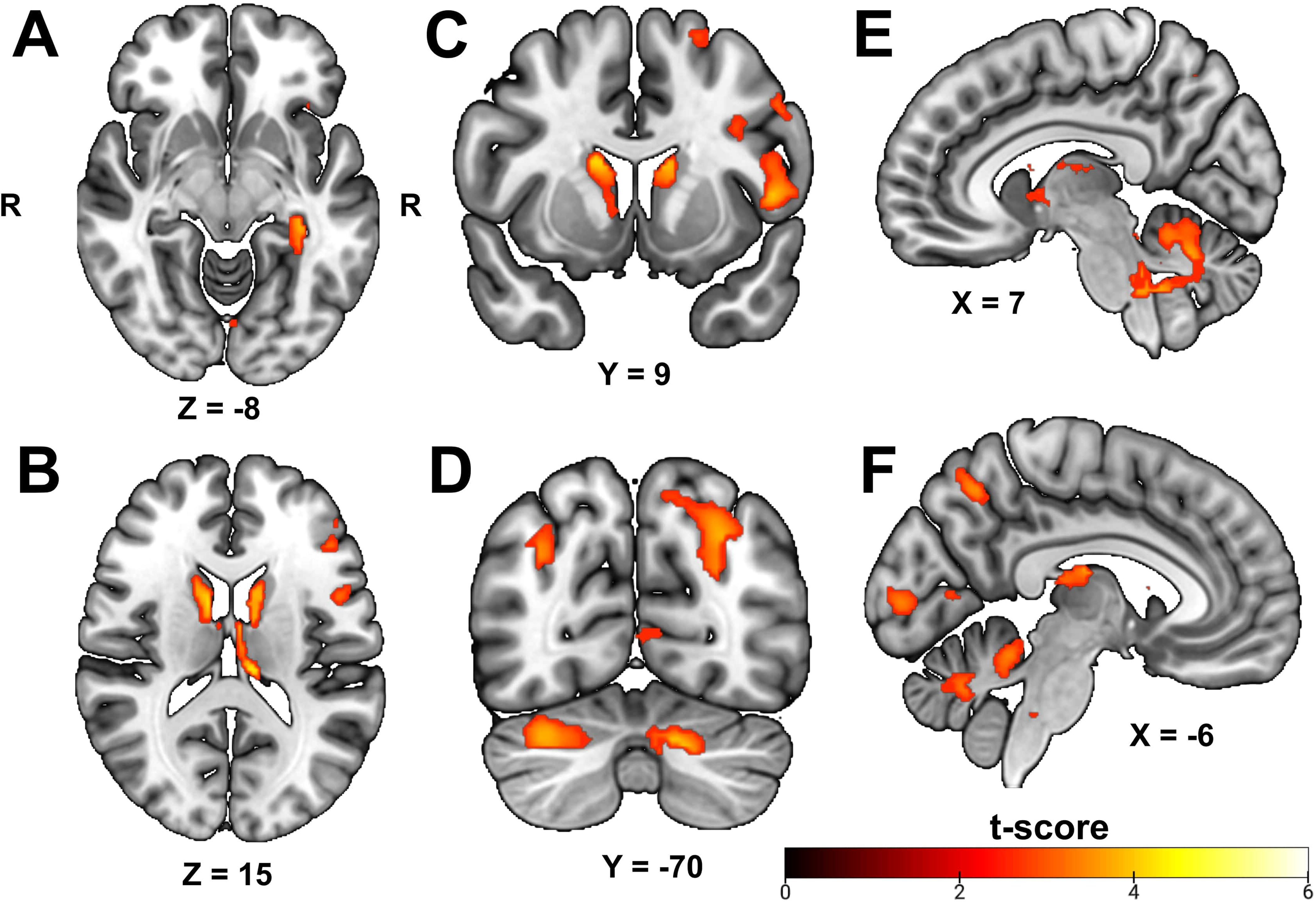
Regional activation positively associated with out-of-scanner long delay recall. Functional magnetic resonance imaging (fMRI) results significantly associated with CVLT long delay recall at p < .05, cluster corrected. For related statistical findings, see Table II. A, Inferior axial perspective. B, Midline axial perspective. C, Anterior coronal perspective. D, Posterior coronal perspective, E, Right medial sagittal perspective. F, Left medial sagittal perspective.

Activation of the right cerebellum, right supramarginal gyrus, left inferior and middle frontal gyri, left precentral gyrus, left cuneus, left precuneus, and bilateral medial frontal gyrus was associated with the combined semantic clustering score across learning and delayed recall trials. Stronger activation was associated with better semantic clustering in all significant areas, except the right supramarginal gyrus which was negatively associated (Figure 4).

**Figure 4.**
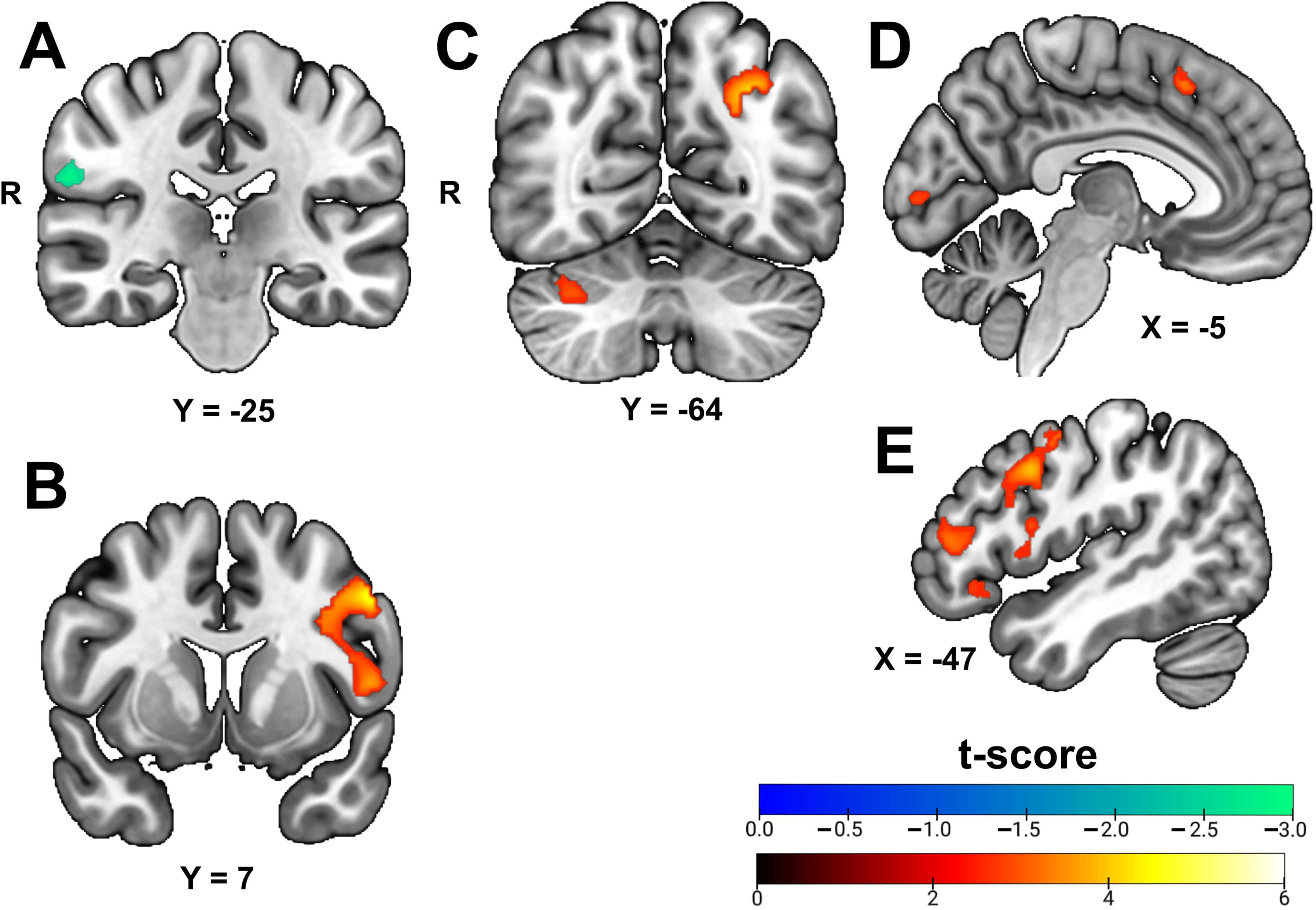
Regional activation associated with semantic clustering. Functional magnetic resonance imaging (fMRI) results significantly associated with CVLT semantic clustering at p < .05, cluster corrected. For related statistical findings, see Table II. A, Anterior coronal perspective. B, Midline coronal perspective. C, Posterior coronal perspective. D, Left medial sagittal perspective, E, Left lateral sagittal perspective (cortical surface).

#### 3.3.2. In-Scanner Experimental Recognition Accuracy

In-scanner experimental recognition accuracy was associated with BOLD activation of 13 regions during encoding (Table II). Experimental accuracy was associated with activity of the: right hippocampus, middle and superior occipital gyri, and supramarginal gyrus; left inferior, middle, medial, and superior frontal gyri, cuneus, and superior parietal lobule; and bilateral caudate and thalamus (Figure 5). A large cluster (>8000 voxels) comprising the right cerebellum and multiple temporal lobe regions (e.g., left hippocampus, left parahippocampal gyrus) was also associated with experimental activity (Figure 6). Greater activity was associated with better experimental recognition accuracy in all regions, except the right supramarginal gyrus.

**Figure 5.**
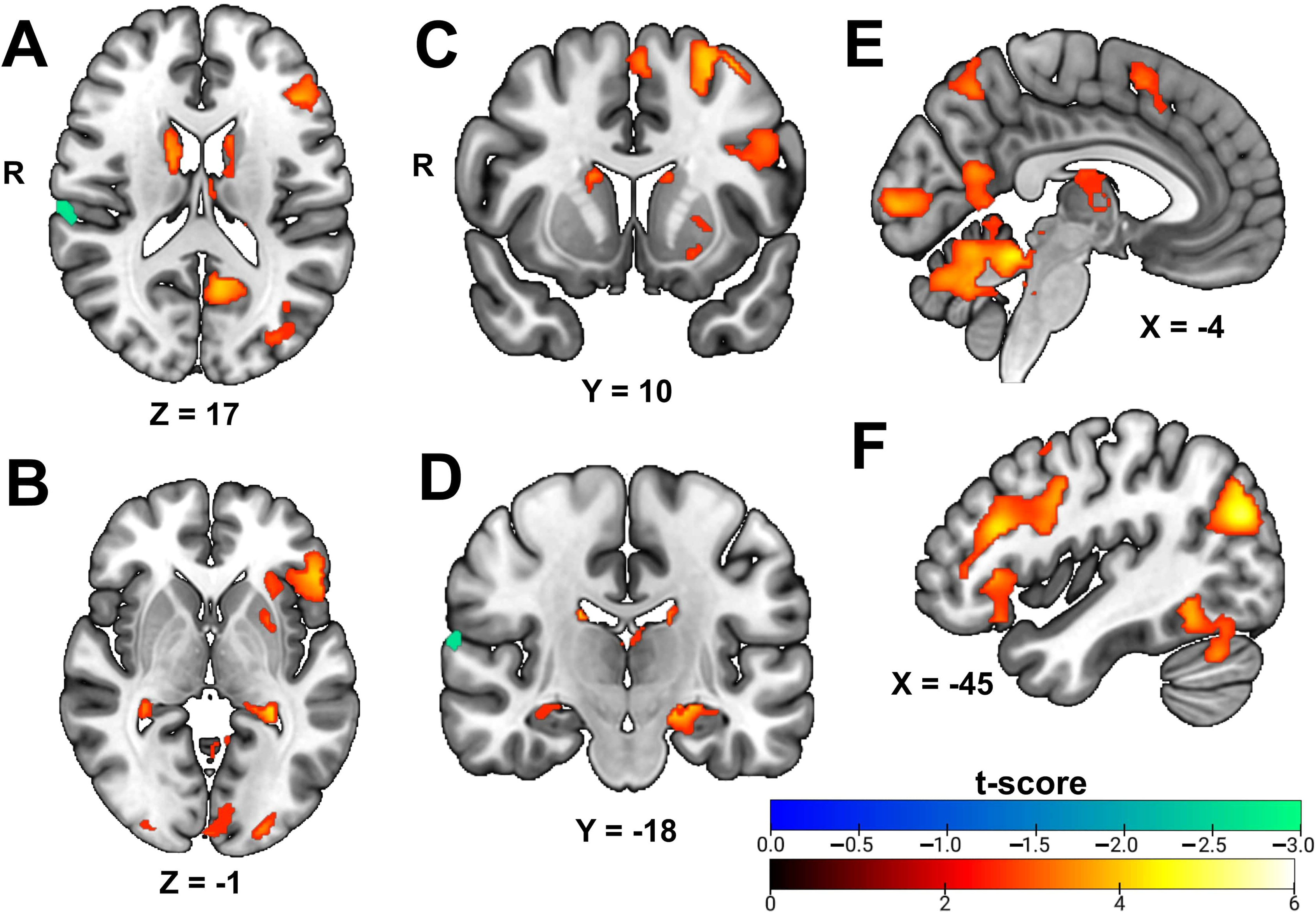
Regional activation associated with in-scanner recognition accuracy. Functional magnetic resonance imaging (fMRI) results significantly associated with in-scanner experimental recognition accuracy at p < .05, cluster corrected. For related statistical findings, see Table II. A, Inferior axial perspective. B, Midline axial perspective. C, Midline coronal perspective. D, Posterior coronal perspective, E, Left medial sagittal perspective. F, Left lateral sagittal perspective (cortical surface).

**Figure 6.**
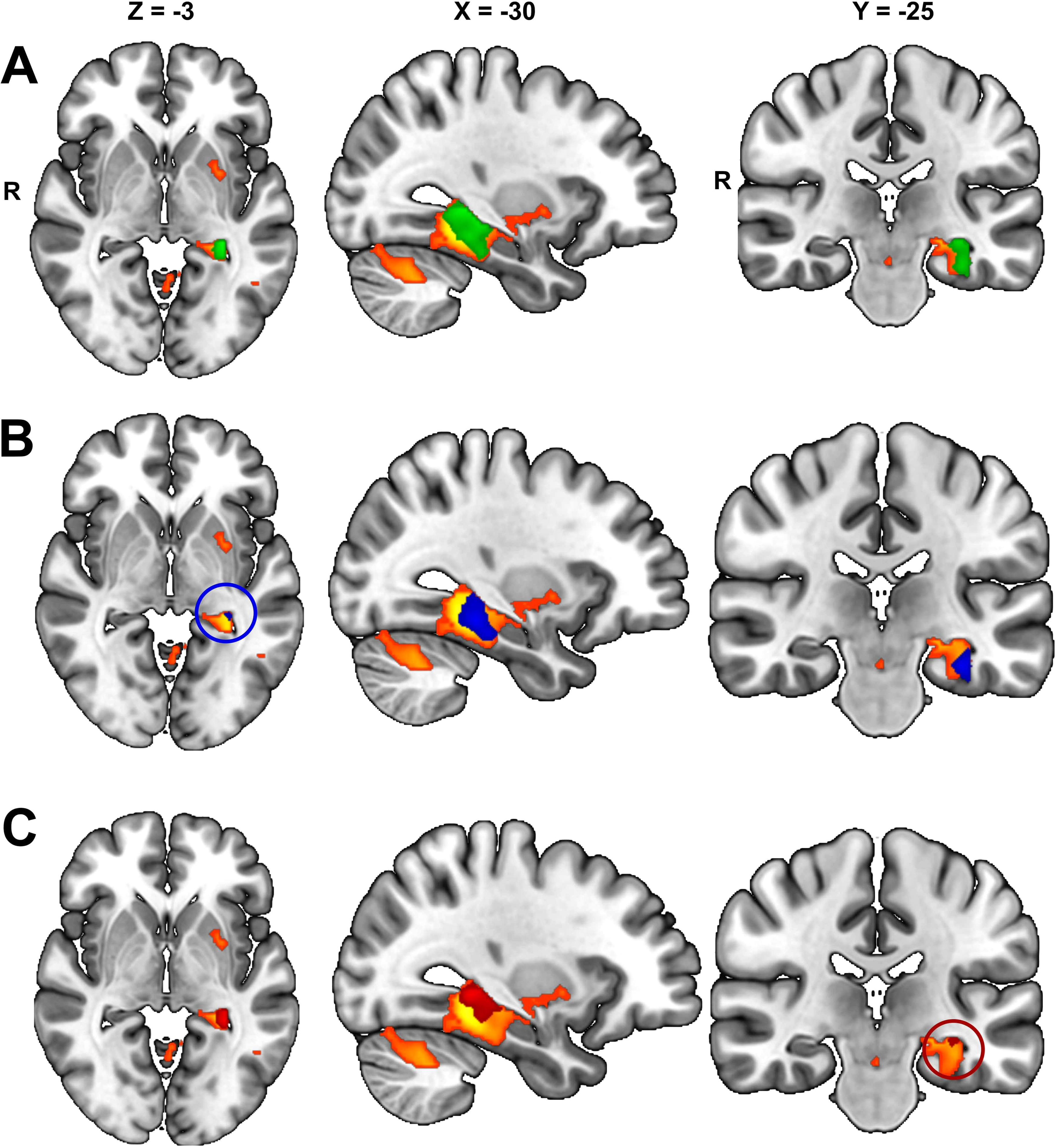
Overlay of large cluster associated with experimental accuracy and other significant clusters. Significant clusters from other analyses superimposed on the large cluster (>8000 voxels; orange) associated with experimental accuracy. For related statistical findings and MNI coordinates of superimposed clusters, see Table II. From left to right: inferior axial perspective, medial sagittal perspective, and posterior coronal perspective. A, Left hippocampus cluster (green) associated with total learning superimposed on the large cluster. B, Left parahippocampal gyrus cluster (blue, circled in axial view) associated with short delay recall superimposed on the large cluster. C, Left hippocampus cluster (red, circled in coronal view) associated with long delay recall superimposed on the large cluster.

Greater brain activation was associated with better performance in all models of verbal learning and memory performance (see Figure 7 for example), except for the associations between right supramarginal gyrus activity and measures of semantic clustering and experimental accuracy which were negatively associated (Figure 8).

**Figure 7.**
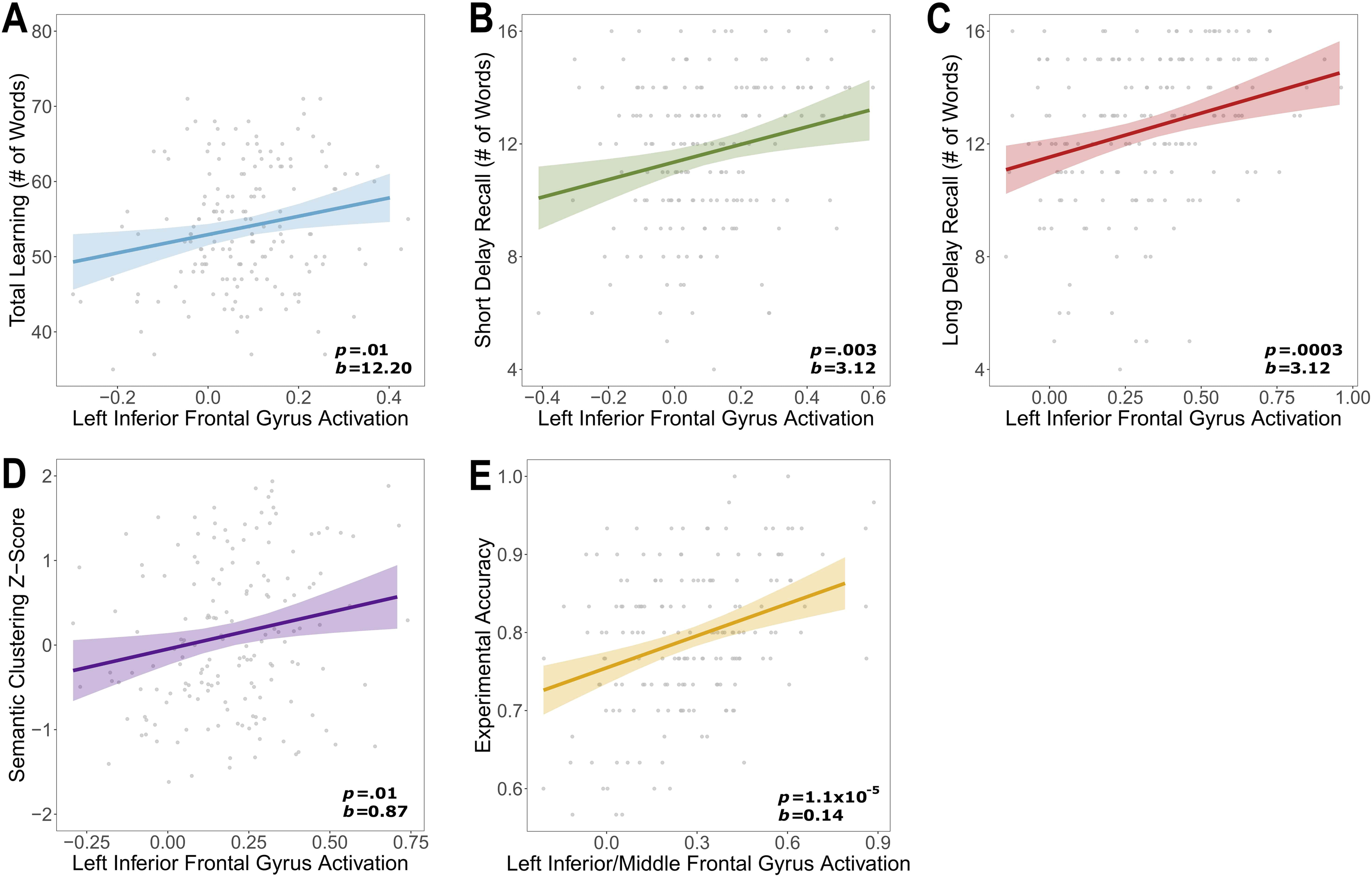
Left inferior frontal gyrus activity by each measure of verbal memory performance. Regression lines demonstrating significant associations between activation of the left inferior frontal gyrus and each measure of verbal memory performance, adjusting for age, race, and education. Individual data points represent raw values. For related statistical findings, see Table II. A-E depict left inferior frontal gyrus activation by: A, total learning score; B, short delay recall; C, long delay recall; D, semantic clustering, E, experimental accuracy.

**Figure 8.**
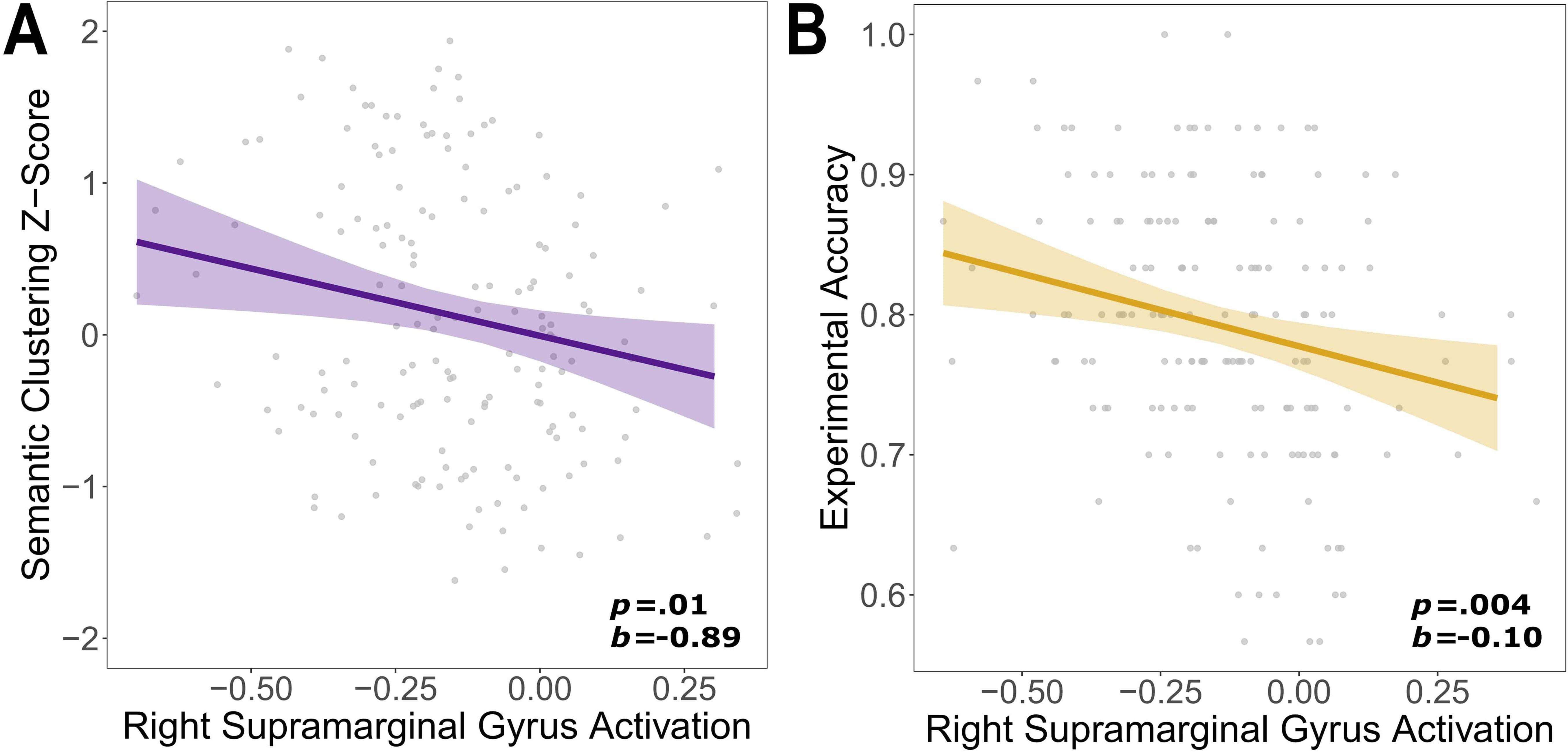
Right supramarginal gyrus activity by semantic clustering and experimental accuracy. Regression lines demonstrating significant associations between activation of the right supramarginal gyrus and measures of semantic clustering and experimental accuracy, adjusting for age, race, and education. Individual data points represent raw values. For related statistical findings, see Table II. A, Semantic clustering by right supramarginal gyrus activity; B, Experimental accuracy by right supramarginal activity.

### 3.4. Functional Connectivity

#### 3.4.1. Left Hippocampal Functional Connectivity

Stronger functional connectivity between the left hippocampus and twelve unique regions was associated with better performance across all CVLT measures (Table III; Figures 9 and 10). Verbal learning was positively associated with stronger connectivity between the left hippocampus and left anterior cingulate gyrus. Short delay recall was positively associated with left hippocampal functional connectivity to the left fusiform and right inferior and middle frontal gyri, while long delay recall was associated with left hippocampal functional connectivity to the left middle cingulate cortex and right medial/superior frontal gyrus. Semantic clustering was associated with stronger connectivity from the left hippocampus to the left fusiform, lingual, and middle cingulate gyri as well as the right middle, medial, and superior frontal gyri. No functional connections with the left hippocampus were associated with in-scanner experimental recognition accuracy.

**Figure 9.**
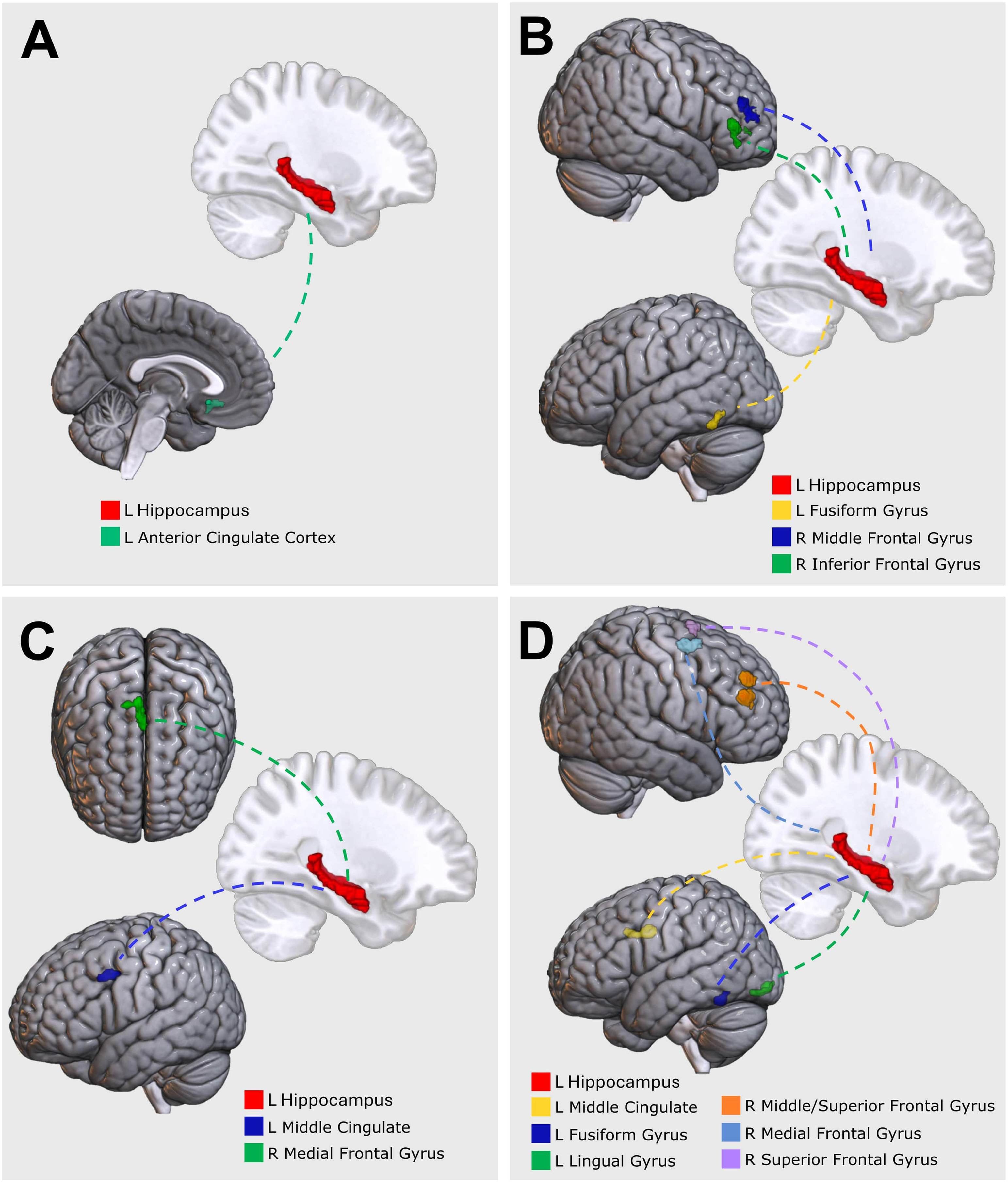
Left hippocampal functional connections for each CVLT measure. For related statistical findings and MNI coordinates, see Table III. Left hippocampal functional connections significantly associated with: A, Total learning; B, Short delay recall; C, Long delay recall; D, Semantic clustering.

**Figure 10.**
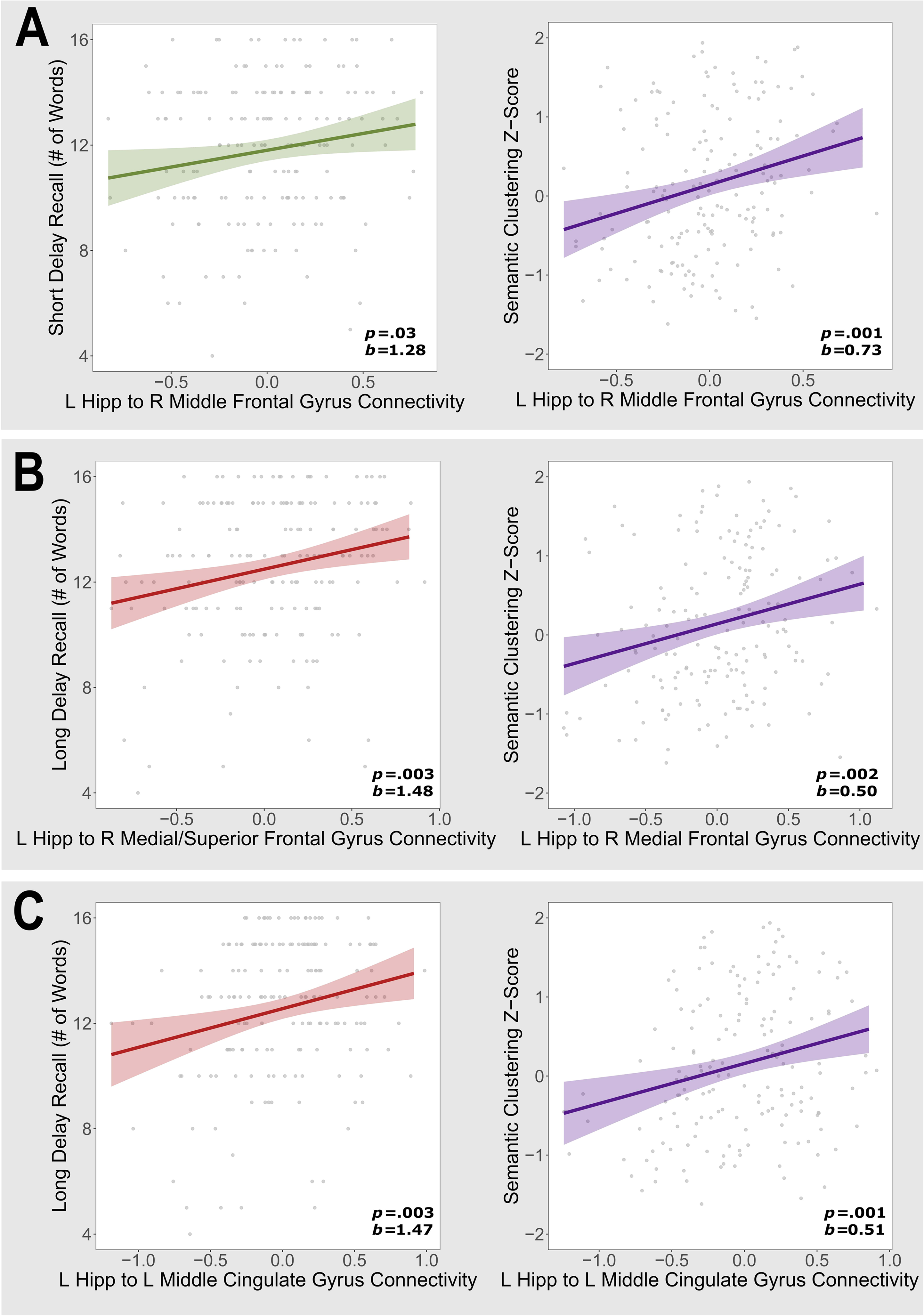
Regression plots of selected left hippocampal functional connectivity findings. Regression lines demonstrating significant associations between CVLT measures and left hippocampal functional connections, adjusting for age, race, and education. Individual data points represent raw values. A, CVLT measures positively associated with functional connectivity between left hippocampus and right middle frontal gyrus; B, CVLT measures positively associated with functional connectivity between left hippocampus and right medial frontal gyrus. C, CVLT measures positively associated with functional connectivity between left hippocampus and left middle cingulate gyrus.

**TABLE III.**
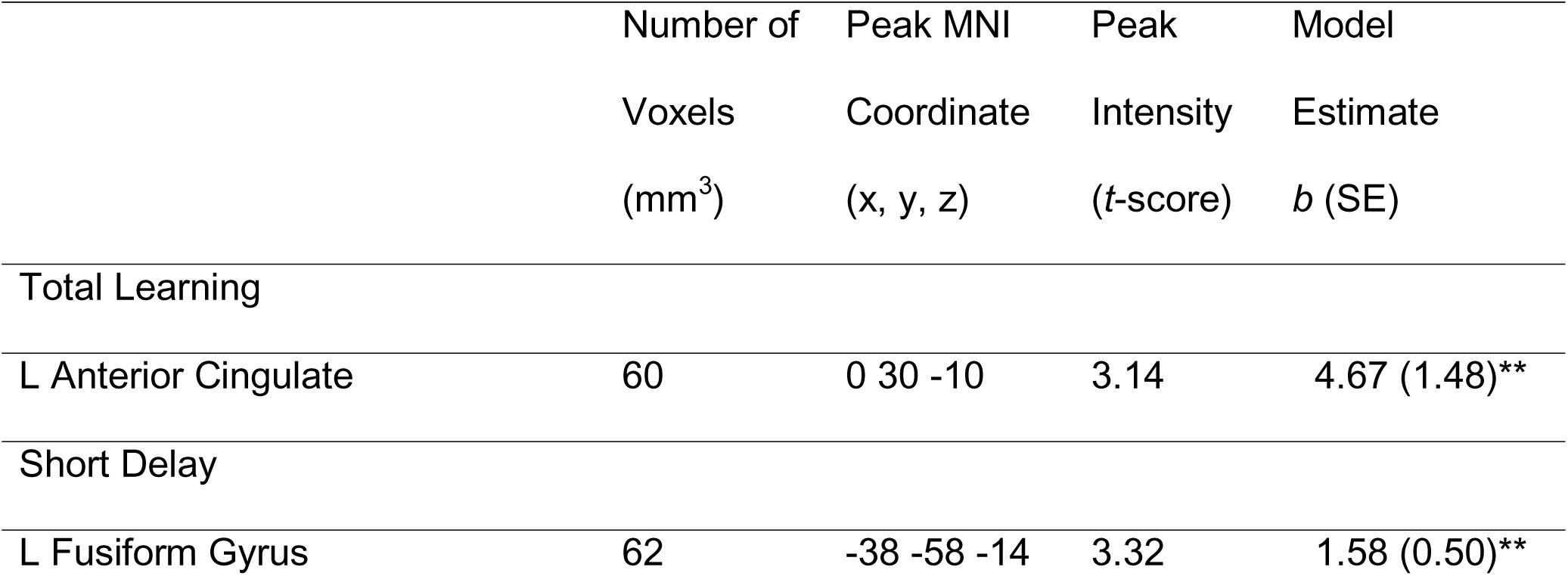
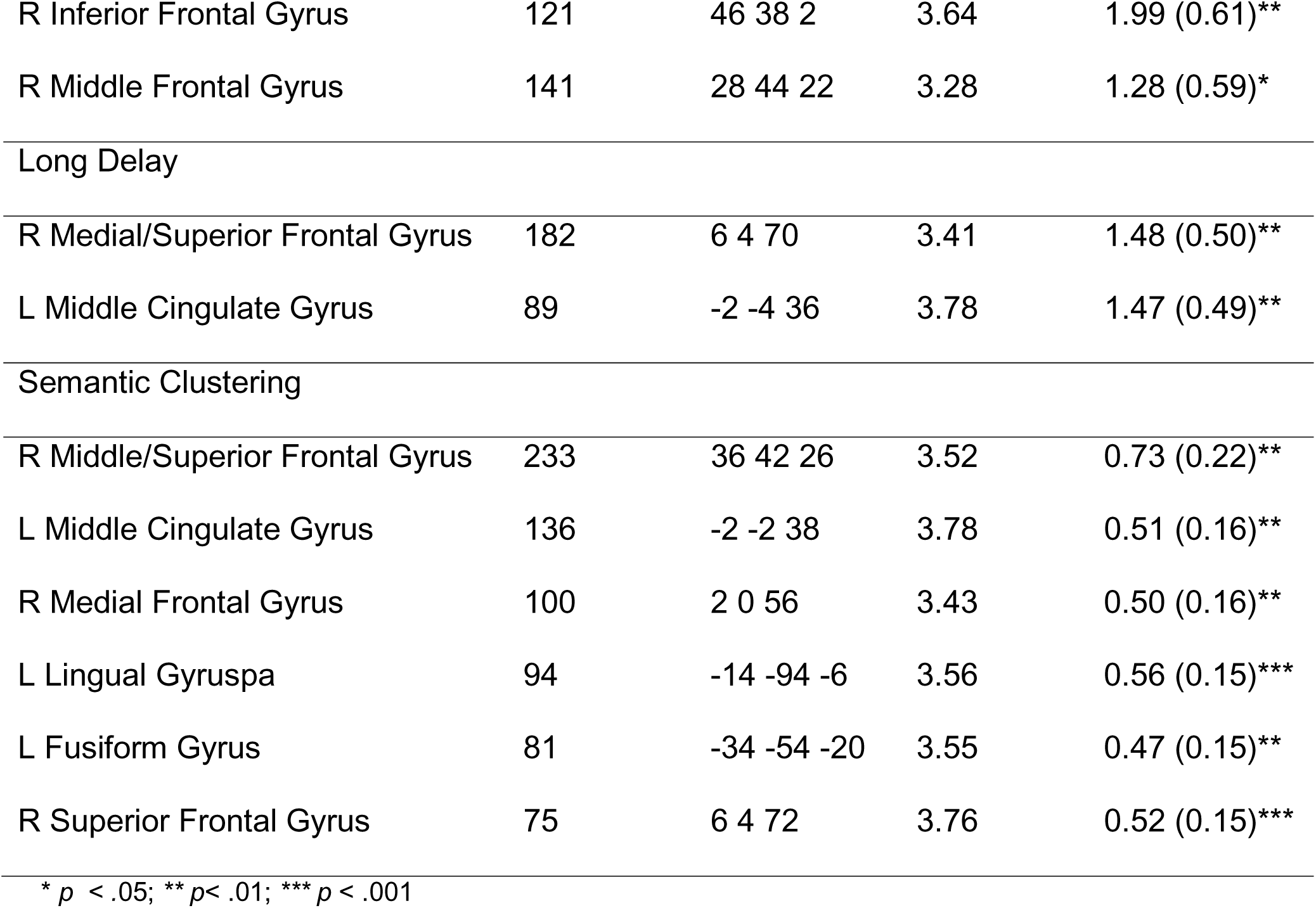
LEFT HIPPOCAMPAL FUNCTIONAL CONNECTIONS ASSOCIATED WITH VERBAL MEMORY MEASURES.

#### 3.4.2. Right Hippocampal Functional Connectivity

Functional connectivity between the right hippocampus and 21 unique regions was associated with CVLT performance (Table IV; Figure 11). Verbal learning was negatively associated with the connectivity from the right hippocampus to the left parahippocampal gyrus (Figure 12) and positively associated with connectivity from the right hippocampus to bilateral medial inferior frontal gyri. Short delay recall was associated with connectivity from the right hippocampus to the right inferior and middle frontal gyri, and to the left middle cingulate gyrus. Long delay recall was associated with right hippocampal functional connectivity to the left middle cingulate gyrus. Semantic clustering was associated with right hippocampal connectivity to the right inferior, medial, middle and superior frontal gyri, and the left middle cingulate gyrus. Finally, in-scanner experimental recognition accuracy was associated with right hippocampal functional connectivity to the left culmen and medial cerebellum, bilateral inferior and middle temporal gyri, and superior parietal lobule (Figure 13). With the exception of the left parahippocampal gyrus which was negatively associated with learning, stronger functional connections from the right hippocampus to all regions was associated with better verbal memory performance (see Figure 14 for examples).

**Figure 11.**
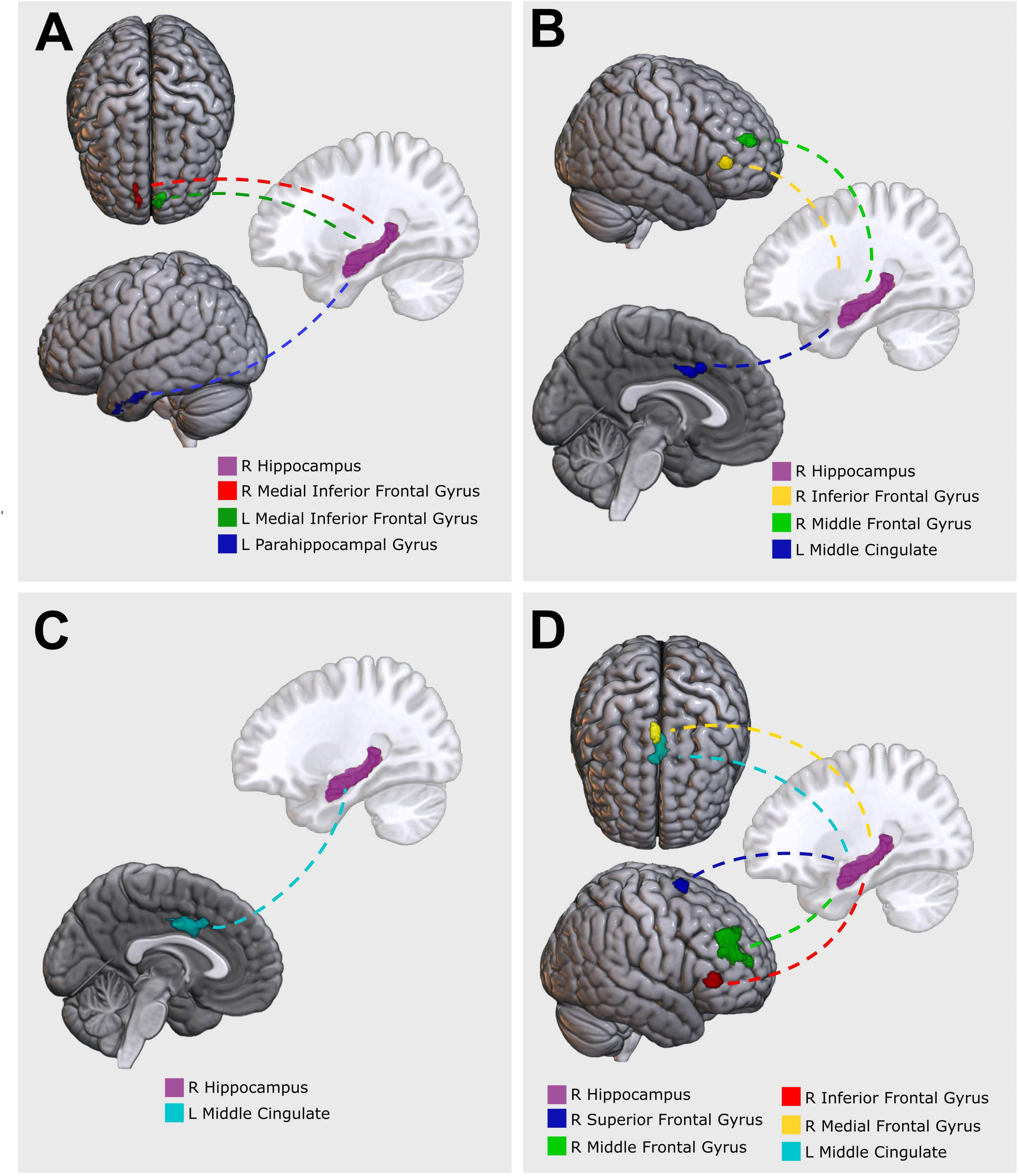
Right hippocampal functional connections for each out-of-scanner verbal memory measure. For related statistical findings and MNI coordinates, see Table IV. Right hippocampal functional connections significantly associated with: A, Total learning; B, Short delay recall; C, Long delay recall; D, Semantic clustering.

**Figure 12.**
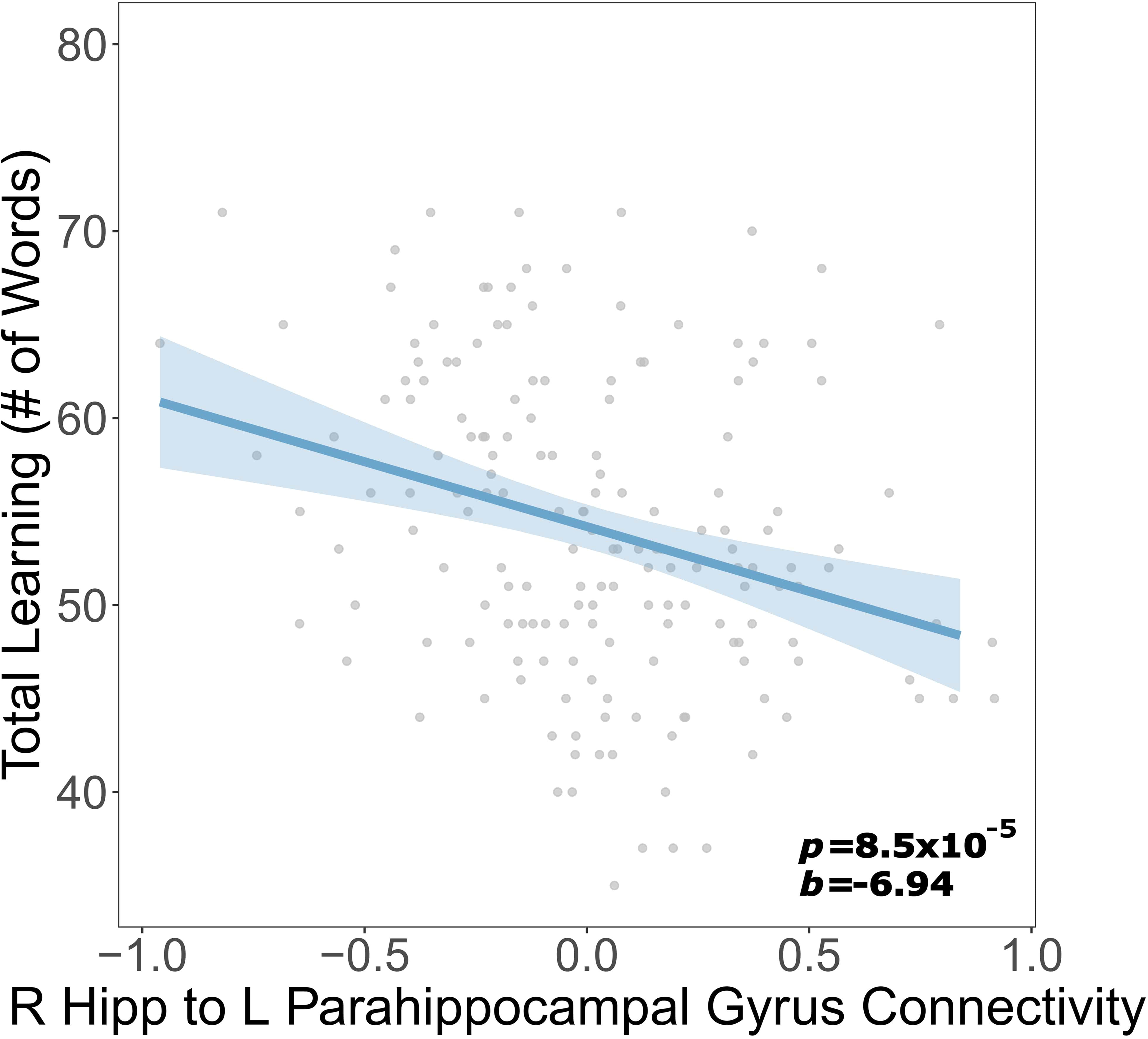
Negative association between total learning score and functional connectivity between the right hippocampus and left parahippocampal gyrus. For related statistical findings and MNI coordinates, see Table IV. Individual data points represent raw scores.

**Figure 13.**
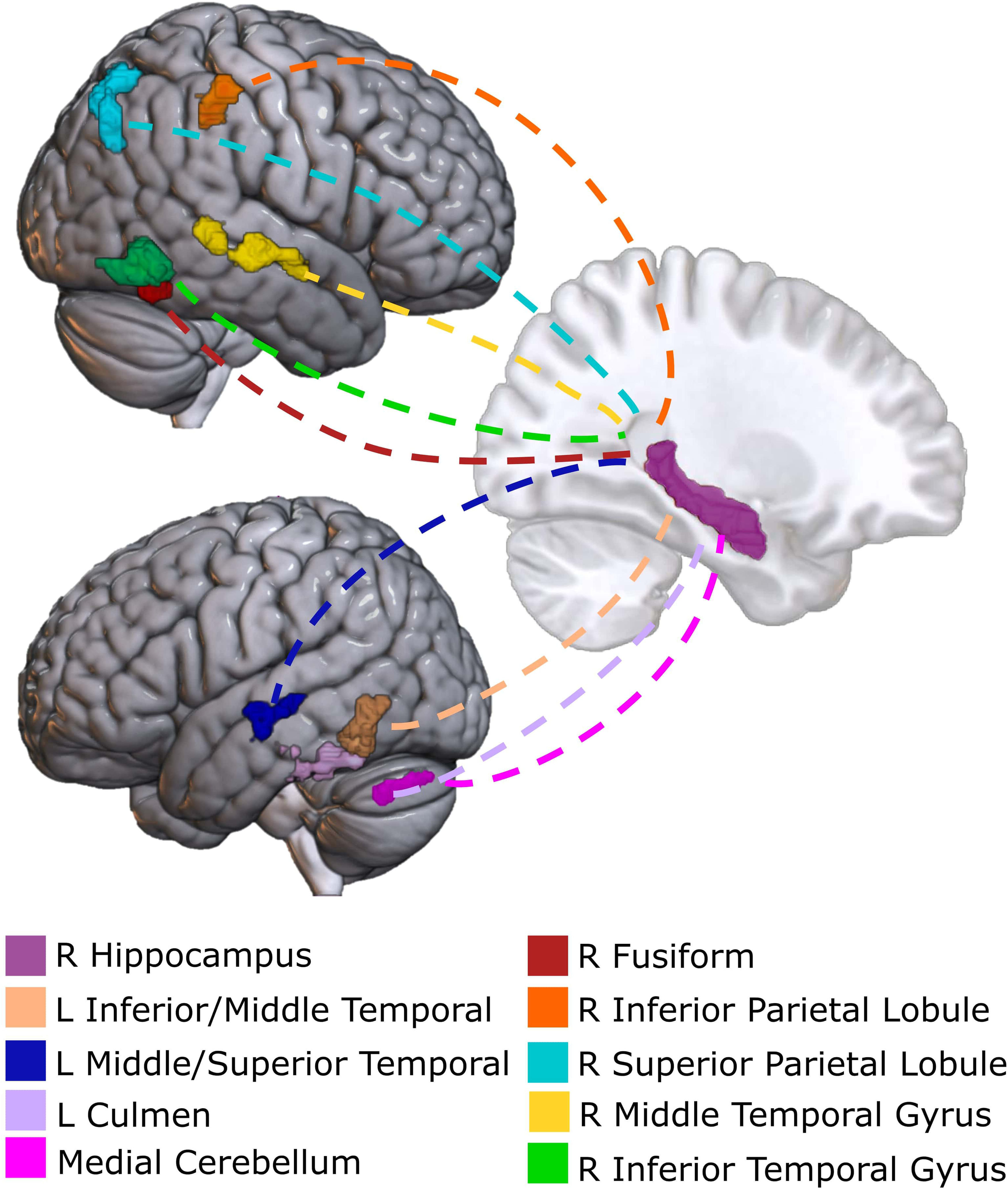
Right hippocampal functional connections associated with in-scanner experimental accuracy. For related statistical findings and MNI coordinates, see Table IV.

**Figure 14.**
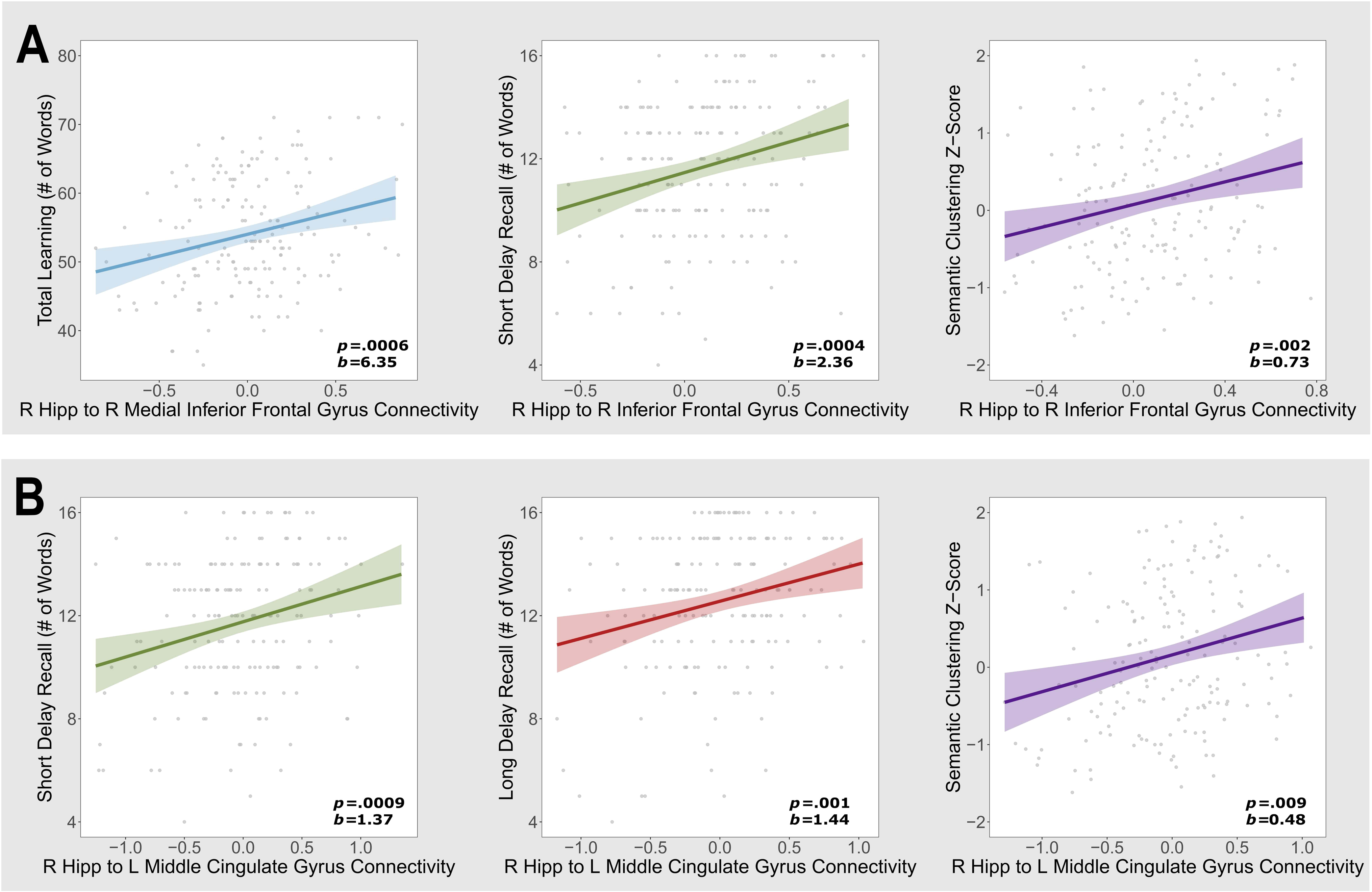
Regression plots of selected right hippocampal functional connectivity findings. Regression lines demonstrating significant associations between CVLT measures and right hippocampal functional connections, adjusting for age, race, and education. Individual data points represent raw scores. For related statistical findings, see Table IV. A, CVLT measures positively associated with functional connectivity between right hippocampus and right inferior frontal gyrus; B, CVLT measures positively associated with functional connectivity between right hippocampus and left middle cingulate gyrus.

**TABLE IV.**
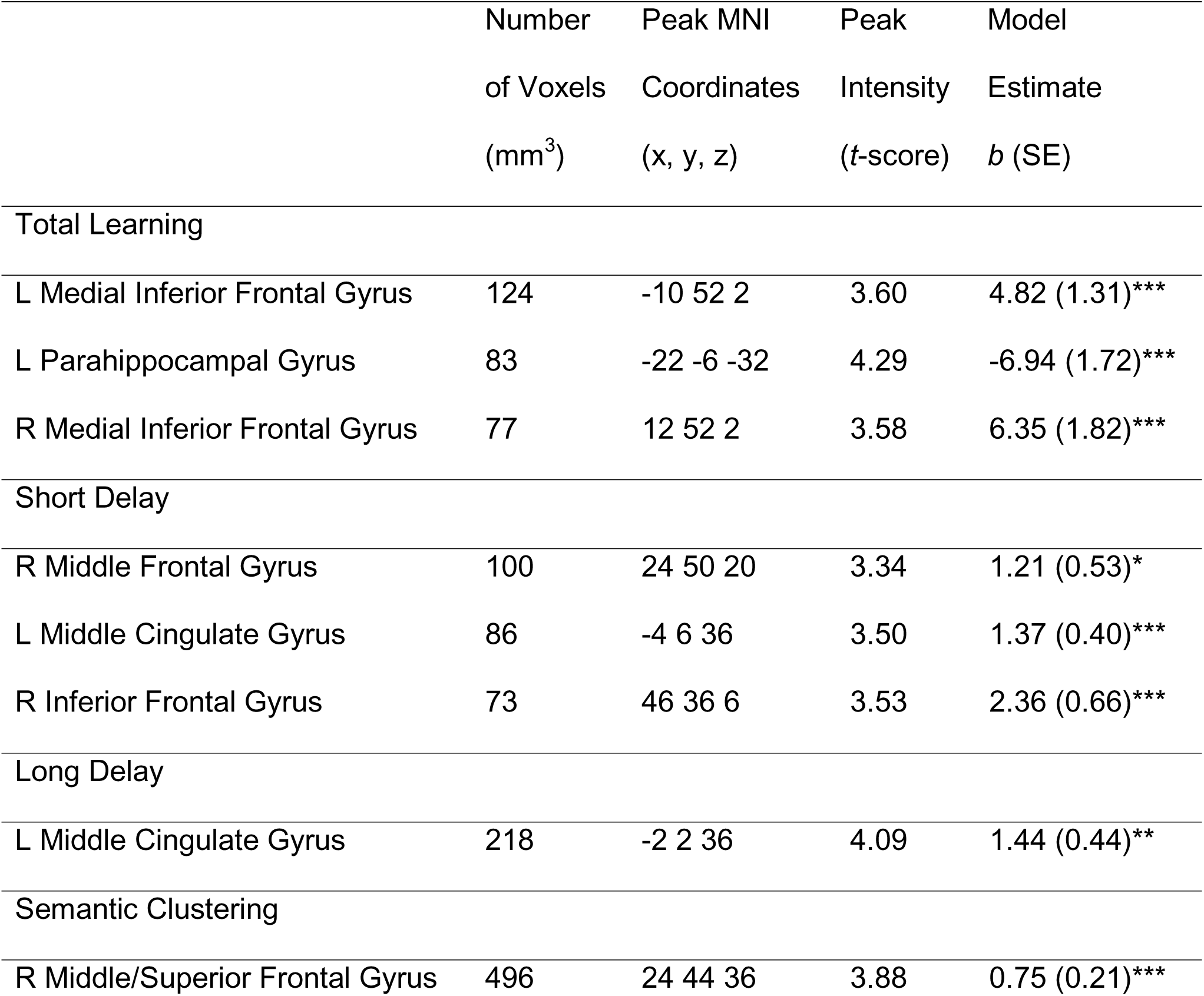
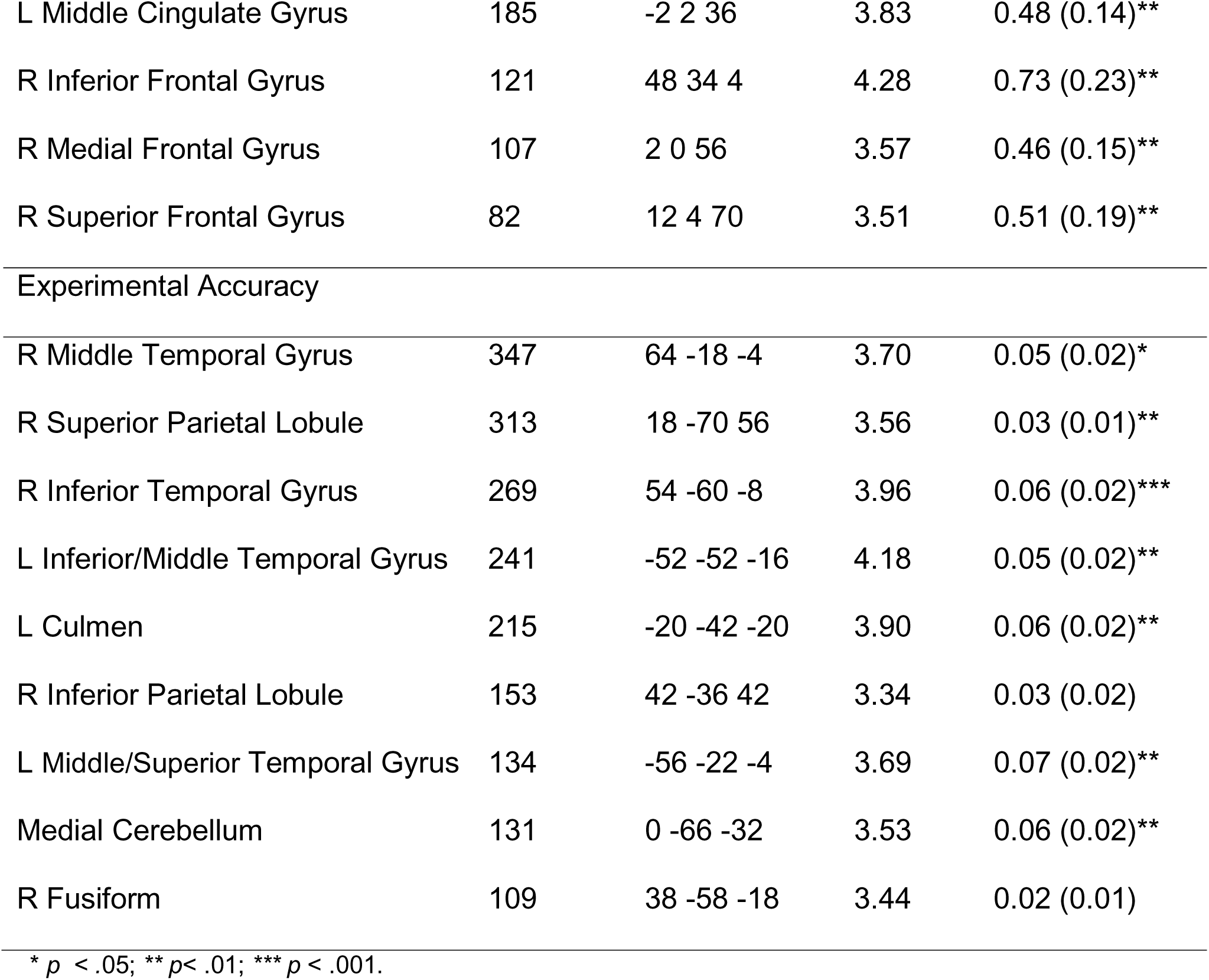
RIGHT HIPPOCAMPAL FUNCTIONAL CONNECTIONS ASSOCIATED WITH VERBAL MEMORY MEASURES.

### 3.5. Supplemental Tertile Analyses

Findings from the supplemental analyses dividing our sample into tertiles by CVLT performance generally reaffirmed the results from the continuous analyses (Appendix A: Supplementary material).

## 4. DISCUSSION

### 4.1. Summary of Findings

In a sample of cognitively normal, midlife postmenopausal women, we investigated the patterns of activation and hippocampal functional connectivity associated with verbal memory performance. Associations between activation and/or connectivity in several brain regions emerged with verbal memory performance. Most notably, greater activation of the left inferior frontal gyrus was associated with better performance on all verbal memory measures. Greater activity of the bilateral medial frontal gyri during encoding was associated with better learning and semantic clustering abilities. Additionally, greater left hippocampal activation was positively associated with multiple verbal memory measures including learning, long delay recall, and in-scanner recognition accuracy. In contrast, regional activation associated with in-scanner experimental recognition accuracy comprised parietal and occipital regions that were not widely associated with CVLT measures. Thus, consistent with prior work in this population (Jacobs et al., 2016; Schroeder et al., 2024), the pattern of activation during verbal encoding that supported successful verbal memory assessed using sensitive list-learning tasks primarily comprised greater activity of left inferior and bilateral medial frontal gyri, as well as the left hippocampus.

Hippocampal functional connectivity analyses revealed findings in multiple brain regions consistent across verbal memory measures. Stronger left and right hippocampal functional connections to regions within the right prefrontal cortex (e.g., right middle and medial frontal gyri) corresponded to better verbal learning, semantic clustering, and delayed recall. Conversely, in-scanner recognition accuracy was primarily associated with right hippocampal functional connections to temporal and parietal regions. Left-to-right hippocampal functional connectivity was not associated with verbal memory performance in this sample; stronger connectivity from the right hippocampus to the left parahippocampal gyrus was the only functional connection associated with worse performance on any verbal memory measure. Overall, these findings suggest that bilateral prefrontal and hippocampal activation, as well as strong functional connections between these regions, support verbal learning and memory in midlife postmenopausal women.

### 4.2. Interpreting Bilateral Prefrontal Activation

In our sample, activations and hippocampal functional connections associated with CVLT measures were largely localized to the PFC. Prefrontal-hippocampal connections are key to episodic encoding (Eichenbaum et al., 2017); however, given past literature indicating the role of the left hemisphere in episodic encoding (see Tulving et al., 1994), the bilaterality of prefrontal activation observed in the present study was somewhat unexpected. According to the Hemispheric Asymmetry Reduction in Older Adults model proposed by Cabeza (2002), older adults utilize less lateralized prefrontal regions for episodic memory processes compared to younger adults. The recruitment of bilateral prefrontal regions in our sample may therefore indicate a shift towards “older” patterns of activation among midlife women. The shift towards bilateral prefrontal regions with aging may be a compensatory mechanism whereby older adults recruit both hemispheres to successfully complete a task that requires only one hemisphere in younger adults (see Cabeza, 2002 for review). Another interpretation is that bilateral prefrontal activation is a result of the dedifferentiation of cognitive domains and an indication of greater reliance on executive functioning with aging (Cabeza, 2002). Additionally, evidence from PET studies points to the left PFC as an important region for semantic clustering (Fletcher et al., 1998; Gabrieli et al., 1998; Savage et al., 2001). Women tend to use semantic strategies more frequently than men (Kramer et al., 1988), and women taking MHT have shown greater semantic clustering compared to non-users (Maki et al., 2001). We previously identified that postmenopausal women with higher endogenous estradiol had greater activation of multiple regions of the PFC (i.e., left inferior frontal gyrus and bilateral middle frontal gyrus) and better learning and semantic clustering scores (Schroeder et al., 2024). Here, we similarly found that greater activation of the left inferior and middle frontal gyri, and bilateral medial frontal gyri was associated with better semantic clustering. Drawing upon our behavioral and neuroimaging findings indicating that women are engaging in semantic clustering, the bilateral pattern of prefrontal activation may reflect compensatory use of effortful semantic learning strategies to sustain verbal memory abilities.

### 4.3. Understanding Deactivation and Negative Functional Connections

It is worth noting that the only negative activations observed were the deactivation of the right supramarginal gyrus, and the negative functional connectivity between the right hippocampus and left parahippocampal gyrus. Previous work in short-term memory has demonstrated that the left supramarginal gyrus is critical for encoding serial order of verbal information independent of semantic content (Guidali et al., 2019). Considering evidence that women less frequently use serial learning strategies in list-learning tasks compared with men (Kramer et al., 1988), the deactivation of the right supramarginal gyrus during verbal encoding in our sample of midlife women may reflect an inhibition of serial learning strategies. However, we interpret this with caution due to the inconsistency in hemispheric laterality between our findings and previous literature (Guidali et al., 2019).

We also found that greater activity of the right hippocampus and simultaneous decreased activity of the left parahippocampal gyrus was associated with better verbal learning. A similar pattern has been noted in two longitudinal studies of MHT use, showing that MHT users had increased hippocampal activity and decreased parahippocampal gyrus activation compared to non-users during verbal recognition (Maki et al., 2011; Resnick et al., 1998). While this may suggest that endogenous estrogen levels are contributing to our findings, we did not previously identify activity of this region to be associated with estrogens in the postmenopause in this sample (Schroeder et al., 2024). Other evidence suggests that deactivation of the parahippocampal gyrus underlies state-related episodic retrieval processes, indicative of engagement in a sustained retrieval mode with ongoing task demands (Donaldson et al., 2001). Given that participants are explicitly instructed to remember the presented words for a later memory test, an alternative explanation is that the parahippocampal gyrus was deactivated relative to the hippocampus because women were engaging in simultaneous recognition processes (e.g., mentally rehearsing items) as an encoding strategy.

### 4.4. Recall vs. Recognition

The variety of verbal memory measures assessed in the present study may have differing memory demands; specifically, the CVLT requires active recollection of learned words, whereas the in-scanner experimental accuracy variable is a recognition measure assessing familiarity of previously viewed words. Recollection typically places greater demands on memory systems than familiarity, likely producing different behavioral results and different neural substrates (for review, see Eichenbaum et al., 2007; Yonelinas, 2002). While average performance scores were similar on recall and recognition paradigms in our sample, differences in patterns of hippocampal functional connectivity supporting recall compared to recognition can be interpreted with evidence from neuroimaging studies of recollection and familiarity (e.g., Henson et al., 1999; Strange et al., 2002). For example, prefrontal and medial temporal activation during encoding is more closely associated with later successful recollection than with familiarity (Henson et al., 1999; Strange et al., 2002). Another explanation for the strong hippocampal-frontal functional connections underlying recall—but not recognition—in our sample is that frontal regions are critical for semantic strategies that facilitate recollection processes (Fletcher et al., 1998; Gabrieli et al., 1998; Savage et al., 2001). The experimental accuracy measure was not conducive to deeper, semantic strategies thus negating the reliance on hippocampal-prefrontal connections underlying this measure. Nevertheless, performance on CVLT measures, in-scanner experimental accuracy, and post-scanner free recall measures were highly correlated in our sample, indicating that no measure was particularly ineffective at capturing participants’ verbal learning and memory abilities relative to the other measures.

### 4.5. Study Design Strengths and Limitations

This is the first study in late midlife women to comprehensively examine the underlying neural circuitry that supports performance on a validated verbal episodic memory task. To our knowledge, this is the largest study assessing both task-based cognitive neuroimaging and neuropsychological outcomes in midlife women. The neuroscience literature has been criticized for its over-reliance on male-only samples and its underuse of optimal designs to assess sex differences, yielding gaps in foundational knowledge in women’s health (Rechlin et al., 2022; Woitowich et al., 2020; Will et al., 2017). Given that our sample comprises cognitively normal, postmenopausal women not taking MHT, we can draw fundamental conclusions regarding the underlying neural circuitry that maintains verbal memory abilities in midlife women. Considering prior work showing sex differences in verbal memory-based neuroimaging outcomes at midlife (Jacobs et al., 2016), we acknowledge that the generalizability of our findings to middle-aged men may be limited. Nevertheless, the present study using a large female-only sample responds to the need for high-quality neuroscience research to improve women’s cognition and brain health outcomes as they age.

The study is limited by some methodological differences between the CVLT and in-scanner memory task. Due to the restraints of the fMRI scanner, oral free recall verbal memory tasks which would be more comparable to the CVLT are difficult to administer during scanning. Furthermore, women may have experienced anxiety, claustrophobia, or difficulty concentrating inside the scanner which may have impacted their memory strategies and abilities on that task more than on the CVLT. Despite these limitations, we reported strong positive correlations between CVLT measures, in-scanner experimental accuracy, and post-scanner free recall in our sample.

### 4.6. Conclusions

We demonstrated that patterns of activation and hippocampal functional connectivity that support verbal learning and memory primarily include bilateral prefrontal and medial temporal regions. Many of these regions are crucial for semantic clustering abilities, indicating that women are utilizing effortful, semantic strategies to facilitate verbal learning and recall. Our findings highlight regions of interest that should be targeted in future pharmacological and lifestyle interventions to help women maintain memory abilities as they age. Finally, these patterns of brain functioning in the postmenopause provide a foundation from which we can better evaluate the effects of hormonal, biological, and affective factors on cognition and brain health among midlife women.

## CRediT authorship contribution statement

**Katrina A. Wugalter:** Conceptualization, Formal analysis, Investigation, Writing – Original draft, Writing – Review & Editing, Visualization. **Rebecca C. Thurston**: Conceptualization, Methodology, Resources, Supervision, Project administration, Funding acquisition, Writing - Review & Editing. **Minjie Wu**: Methodology, Software, Formal analysis, Supervision, Writing - Review & Editing. **Rachel A. Schroeder**: Conceptualization, Writing - Review & Editing. **Howard J. Aizenstein**: Methodology, Validation, Writing - Review & Editing. **Pauline M. Maki**: Conceptualization, Methodology, Resources, Supervision, Project administration, Funding acquisition, Writing - Review & Editing.

## References

Balthazar, M.L.F., Yasuda, C.L., Cendes, F., & Damasceno, B.P. (2010). Learning, retrieval, and recognition are compromised in aMCI and mild AD: Are distinct episodic memory processes mediated by the same anatomical structures? Journal of the International Neuropsychological Society, 16, 205–209.

Behzadi, Y., Restom, K., Liau, J., & Liu, T. T. (2007). A component-based noise correction method (CompCor) for BOLD and perfusion based fMRI. NeuroImage, 37(1), 90–101. 10.1016/j.neuroimage.2007.04.042

Brandt, J., & Benedict, R. (2001). Hopkins verbal learning test-revised: professional manual. Psychological Assessment Resources.

Brinton, R.D., Yao, J., Yin, F., Mack, W.J., & Cadenas, E. (2015). Perimenopause as a neurological transition state. Endocrinology, 11.

Cabeza, R. (2002). Hemispheric asymmetry reduction in older adults: The HAROLD model. Psychology and Aging, 17(1), 85–100. 10.1037/0882-7974.17.1.85

Cox, R. W. (1996). AFNI: Software for Analysis and Visualization of Functional Magnetic Resonance Neuroimages. Computers and Biomedical Research, 29(3), 162–173. 10.1006/cbmr.1996.0014

Delis, D. C., Kramer, J. H., Kaplan, E., & Ober, B. A. (2002). California Verbal Learning Test-Second Edition, Adult Version. The Psychological Corporation., 17(5), 509–512. 10.1093/arclin/17.5.509

Donaldson, D. I., Petersen, S. E., Ollinger, J. M., & Buckner, R. L. (2001). Dissociating State and Item Components of Recognition Memory Using fMRI. NeuroImage, 13(1), 129–142. 10.1006/nimg.2000.0664

Eichenbaum, H. (2017). Prefrontal-hippocampal interactions in episodic memory. Nature Reviews Neuroscience, 18, 547–558. 10.1038/nrn.2017.74

Eichenbaum, H., Yonelinas, A. P., & Ranganath, C. (2007). The Medial Temporal Lobe and Recognition Memory. Annual Review of Neuroscience, 30(1), 123–152. 10.1146/annurev.neuro.30.051606.094328

Estevez-Gonzalez, A., Kulisevsky, J., Boltes, A., Otermin, P., & Garcia-Sanchez, C. (2003). Rey verbal learning test is a useful for differential diagnosis in the preclinical phase of Alzheimer’s disease: comparison with mild cognitive impairment and normal aging. International Journal of Geriatric Psychiatry, 18, 1021–1028. 10.1002/gps.1010

Epperson, C.N., Sammel, M.D., & Freeman, E.W. (2013). Menopause effects on verbal memory: findings from a longitudinal community cohort. Journal of Clinical Endocrinology & Metabolism, 98(9), 3829–3838.

Fletcher, P., Shallice, T., & Dolan, R.J. (1998). The functional roles of prefrontal cortex in episodic memory. I. Encoding. Brain, 121(7), 1239–1248. 10.1093/brain/121.7.1239

Friston, K.J., Buechel, C., Fink, G.R., Morris, J., Rolls, E., & Dolan, R.J. (1997). Psychophysiological and modulatory interactions in neuroimaging. Neuroimage, 6(3), 218–229. 10.1006/nimg.1997.0291

Gabrieli, J. D. E., Poldrack, R. A., & Desmond, J. E. (1998). The role of left prefrontal cortex in language and memory. Proceedings of the National Academy of Sciences, 95(3), 906–913. 10.1073/pnas.95.3.906

Gold, E.B., Sternfeld, B., Kelsey, J.L., Brown, C., Mouton, C., Reame, N., Salamone, L., & Stellato, R. (2000). Relation of demographic and lifestyle factors to symptoms in a multi-racial/ethnic population of women 40-55 years of age. American Journal of Epidemiology, 152(5), 463–473.

Grady, C. L., Springer, M. V., Hongwanishkul, D., McIntosh, A. R., & Winocur, G. (2006). Age-related changes in brain activity across the adult lifespan. Journal of Cognitive Neuroscience, 18(2), 227–241. 10.1162/089892906775783705

Greendale, G. A., Huang, M. H., Wight, R. G., Seeman, T., Luetters, C., Avis, N. E., Johnston, J., & Karlamangla, A. S. (2009). Effects of the menopause transition and hormone use on cognitive performance in midlife women. Neurology, 72(21), 1850–1857. 10.1212/WNL.0b013e3181a71193

Guidali, G., Pisoni, A., Bolognini, N., & Papagno, C. (2019). Keeping order in the brain: The supramarginal gyrus and serial order in short-term memory. Cortex, 119, 89–99. 10.1016/j.cortex.2019.04.009

Henson, R. N. A., Rugg, M. D., Shallice, T., Josephs, O., & Dolan, R. J. (1999). Recollection and Familiarity in Recognition Memory: An Event-Related Functional Magnetic Resonance Imaging Study. The Journal of Neuroscience, 19(10), 3962–3972. 10.1523/JNEUROSCI.19-10-03962.1999

Jacobs, E. G., Weiss, B. K., Makris, N., Whitfield-Gabrieli, S., Buka, S. L., Klibanski, A., & Goldstein, J. M. (2016). Impact of sex and menopausal status on episodic memory circuitry in early midlife. The Journal of Neuroscience, 36(39), 10163–10173. 10.1523/JNEUROSCI.0951-16.2016

Kilpi, F., Soares, A. L. G., Fraser, A., Nelson, S. M., Sattar, N., Fallon, S. J., Tilling, K., & Lawlor, D. A. (2020). Changes in six domains of cognitive function with reproductive and chronological ageing and sex hormones: a longitudinal study in 2411 UK mid-life women. BMC Women’s Health, 20(1), 177. 10.1186/s12905-020-01040-3

Kramer, J. H., Delis, D. C., & Daniel, M. (1988). Sex differences in verbal learning. Journal of Clinical Psychology, 44(6), 907–915. 10.1002/1097-4679(198811)44:6<907::AID-JCLP2270440610>3.0.CO;2-8

Kwon, D., Maillet, D., Pasvanis, S., Ankudowich, E., Grady, C. L., & Rajah, M. N. (2016). Context Memory Decline in Middle Aged Adults is Related to Changes in Prefrontal Cortex Function. Cerebral Cortex, 26 (6), 2440–2460. 10.1093/cercor/bhv068

Lange, K. L., Bondi, M. W., Salmon, D. P., Galasko, D., Delis, D. C., Thomas, R. G., & Thal, L. J. (2002). Decline in verbal memory during preclinical Alzheimer’s disease: examination of the effect of APOE genotype. Journal of the International Neuropsychological Society, 8(7), 943–955. 10.1017/s1355617702870096

Maki, P. M., Dennerstein, L., Clark, M., Guthrie, J., LaMontagne, P., Fornelli, D., Little, D., Henderson, V. W., & Resnick, S. M. (2011). Perimenopausal use of hormone therapy is associated with enhanced memory and hippocampal function later in life. Brain Research, 1379, 232–243. 10.1016/j.brainres.2010.11.030

Maki, P.M., Springer, G., Anastos, K., Gustafson, D.R., Weber, K., Vance, D., Dykxhoorn, D., Milam, J., Adimora, A.A., Kassaye, S.G., Waldrop, D., & Rubin, L. (2021). Cognitive changes during the menopausal transition: a longitudinal study in women with and without HIV. Menopause, 28(4), 1–9. 10.1097/gme.0000000000001725

Maki, P. M., Zonderman, A. B., & Resnick, S. M. (2001). Enhanced Verbal Memory in Nondemented Elderly Women Receiving Hormone-Replacement Therapy. American Journal of Psychiatry, 158(2), 227–233. 10.1176/appi.ajp.158.2.227

Milani, S. A., Marsiske, M., Cottler, L. B., Chen, X., & Striley, C. W. (2018). Optimal cutoffs for the Montreal Cognitive Assessment vary by race and ethnicity. Alzheimer’s & Dementia, 10, 773–781. 10.1016/j.dadm.2018.09.003

Nasreddine, Z. S., Phillips, N. A., Bédirian, V., Charbonneau, S., Whitehead, V., Collin, I., Cummings, J. L., & Chertkow, H. (2005). The Montreal Cognitive Assessment, MoCA: a brief screening tool for mild cognitive impairment. Journal of the American Geriatrics Society, 53(4), 695–699. 10.1111/j.1532-5415.2005.53221.x

R Core Team (2023). R: A Language and Environment for Statistical Computing. R Foundation for Statistical Computing, Vienna, Austria. https://www.R-project.org/.

Rechlin, R. K., Splinter, T. F. L., Hodges, T. E., Albert, A. Y., & Galea, L. A. M. (2022). An analysis of neuroscience and psychiatry papers published from 2009 and 2019 outlines opportunities for increasing discovery of sex differences. Nature Communications, 13(1), 2137. 10.1038/s41467-022-29903-3

Resnick, S. M., Maki, P. M., Golski, S., Kraut, M. A., & Zonderman, A. B. (1998). Effects of estrogen replacement therapy on PET cerebral blood flow and neuropsychological performance. Hormones and behavior, 34(2), 171–182. 10.1006/hbeh.1998.1476

Savage, C. R., Deckersbach, T., Heckers, S., Wagner, A. D., Schacter, D. L., Alpert, N. M., Fischman, A. J., & Rauch, S. L. (2001). Prefrontal regions supporting spontaneous and directed application of verbal learning strategies: Evidence from PET. Brain, 124(1), 219–231. 10.1093/brain/124.1.219

Schroeder, R.A., Thurston, R.C., Wu, M., Aizenstein, H.J., Derby, C.A., & Maki, P.M. (2024). Endogenous estrogens and brain activation during verbal memory encoding and recognition in the postmenopause. The Journal of Clinical Endocrinology & Metabolism, 00, 1–10. 10.1210/clinem/dgae467

Strange, B. A., Otten, L. J., Josephs, O., Rugg, M. D., & Dolan, R. J. (2002). Dissociable Human Perirhinal, Hippocampal, and Parahippocampal Roles during Verbal Encoding. The Journal of Neuroscience, 22(2), 523–528. 10.1523/JNEUROSCI.22-02-00523.2002

Thurston, R. C., Chang, Y., Barinas-Mitchell, E., Jennings, J. R., Landsittel, D. P., Santoro, N., Von Känel, R., & Matthews, K. A. (2016). Menopausal Hot Flashes and Carotid Intima Media Thickness Among Midlife Women. Stroke, 47(12), 2910–2915. 10.1161/STROKEAHA.116.014674

Thurston, R. C., Wu, M., Chang, Y.-F., Aizenstein, H. J., Derby, C. A., Barinas-Mitchell, E. A., & Maki, P. (2023). Menopausal Vasomotor Symptoms and White Matter Hyperintensities in Midlife Women. Neurology, 100(2). 10.1212/WNL.0000000000201401

Tulving, E., Kapur, S., Craik, F.I.M., Moscovitch, M., & Houle, S. (1994). Hemispheric encoding/retrieval asymmetry in episodic memory: Positron emission tomography findings. PNAS, 91, 2016-2000.

Tzourio-Mazoyer, N., Landeau, B., Papathanassiou, D., Crivello, F., Etard, O., Delcroix, N., Mazoyer, B., & Joliot, M. (2002). Automated Anatomical Labeling of Activations in SPM Using a Macroscopic Anatomical Parcellation of the MNI MRI Single-Subject Brain. NeuroImage, 15(1), 273–289. 10.1006/nimg.2001.0978

Whitfield-Gabrieli, S., & Nieto-Castanon, A. (2012). Conn: a functional connectivity toolbox for correlated and anticorrelated brain networks. Brain connectivity, 2(3), 125–141. 10.1089/brain.2012.0073

Will, T. R., Proaño, S. B., Thomas, A. M., Kunz, L. M., Thompson, K. C., Ginnari, L. A., Jones, C. H., Lucas, S.-C., Reavis, E. M., Dorris, D. M., & Meitzen, J. (2017). Problems and Progress regarding Sex Bias and Omission in Neuroscience Research. Eneuro, 4(6), ENEURO.0278-17.2017. 10.1523/ENEURO.0278-17.2017

Woitowich, N. C., Beery, A., & Woodruff, T. (2020). A 10-year follow-up study of sex inclusion in the biological sciences. eLife, 9, e56344. 10.7554/eLife.56344

Yonelinas, A. P. (2002). The Nature of Recollection and Familiarity: A Review of 30 Years of Research. Journal of Memory and Language, 46(3), 441–517. 10.1006/jmla.2002.2864

