## Supplemental Table 1 for "Patterns of Brain Activation and Hippocampal Functional Connectivity Supporting Verbal Memory in Midlife Women"

**TABLE S1**

PAIRWISE TERTILE GROUP COMPARISON ESTIMATES

|  | Model Estimate  *b* (SE) | | |
| --- | --- | --- | --- |
|  | High vs. Low | High vs. Middle | Middle vs. Low |
| Total Learning |  |  |  |
| L Hippocampus | 0.10 (0.021)*** | 0.11 (0.022)*** | 0.01 (0.021) |
| L Inferior Frontal Gyrus | 0.07 (0.025)* | 0.06 (0.026)* | 0.01 (0.024) |
| R Hippocampus | 0.08 (0.022)** | 0.05 (0.023) | 0.03 (0.021) |
| R Inferior Frontal Gyrus | 0.06 (0.025)* | 0.05 (0.025) | 0.02 (0.024) |
| Short Delay Recall |  |  |  |
| L Inferior Frontal Gyrus | 0.10 (0.037)* | 0.08 (0.037) | 0.03 (0.035) |
| R Inferior Frontal Gyrus | 0.04 (0.023) | 0.001 (0.023) | 0.04 (0.022) |
| L Hipp to R Inferior Frontal Gyrus | 0.19 (0.063)** | 0.13 (0.063) | 0.06 (0.061) |
| R Hipp to R Inferior Frontal Gyrus | 0.18 (0.059)** | 0.14 (0.059)* | 0.04 (0.055) |
| Long Delay Recall |  |  |  |
| L Hippocampus | 0.08 (0.024)** | 0.04 (0.023) | 0.04 (0.022) |
| L Inferior Frontal Gyrus | 0.14 (0.046)** | 0.05 (0.043) | 0.09 (0.042) |
| Semantic Clustering |  |  |  |
| L Inferior Frontal Gyrus | 0.09 (0.038)* | 0.03 (0.037) | 0.06 (0.038) |
| L Inferior/Middle Frontal Gyrus | 0.13 (0.042)** | 0.10 (0.041) | 0.03 (0.041) |
| R Hipp to R Inferior Frontal Gyrus | 0.14 (0.058)* | 0.07 (0.057) | 0.07 (0.057) |

*Note:* The sample was divided into tertiles (high, medium, low) by performance on each CVLT measure. * *p < .*05; *** p*< .01; **** p* < .001.
